# Selective loss of *Nkx2.1*-lineage neurons in the lateral septum alters the balance between novelty seeking and threat avoidance

**DOI:** 10.1101/2023.09.24.559205

**Authors:** Miguel Turrero García, Diana N. Tran, Ralph E. Peterson, Leena A. Ibrahim, Christopher M. Reid, Sarah M. Hanson, Yuqi Ren, Fiona Dale-Huang, Yajun Xie, Sarah K. Stegmann, Steve Vu, Corey C. Harwell

**Author notes:** Department of Biological Sciences, Texas Tech University. BioScience Program, BESE, King Abdullah University of Science and Technology, Saudi Arabia. Authors for correspondence (M.T.G.), (C.C.H.).

## Abstract

When interacting with their environment, animals must balance exploratory and defensive behavior to evaluate and respond to potential threats. The lateral septum (LS) is a structure in the ventral forebrain that calibrates the magnitude of behavioral responses to stress-related external stimuli, including the regulation of threat avoidance. The complex connectivity between the LS and other parts of the brain, together with its largely unexplored neuronal diversity, makes it difficult to understand how defined LS circuits control specific behaviors. Here, we describe a mouse model where the deletion of the transcriptional regulator *Prdm16* in cells with a common developmental origin (*Nkx2.1*-lineage) results in the almost complete ablation of neurons from this lineage in the LS. Using a combination of single-nucleus RNA sequencing, histological and electrophysiological methods and behavioral analyses, we discovered that *Crhr2*-expresssing neurons are specifically affected in mutant mice, resulting in connectivity and electrophyisiological defects. This neuronal population is specifically activated in stressful contexts, and its removal results in increased exploratory behavior, even under stressful conditions. Our study extends the current knowledge about how defined neuronal populations within the LS can evaluate contextual information to select appropriate behavioral responses. This is a necessary step towards understanding the crucial role that the LS plays in neuropsychiatric conditions where defensive behavior is dysregulated, such as anxiety and aggression disorders.

## INTRODUCTION

The septum is a structure located in the mammalian ventral forebrain, between the lateral ventricles. It is subdivided into medial and lateral nuclei, each with distinct connectivity patterns and functions. The lateral septum (LS) controls different aspects of affective and motivated behavior through its complex connections to and from multiple other brain areas^1–4^. The LS has been classically subdivided into three nuclei (dorsal [LSd], intermediate [LSi], and ventral [LSv]), whose specific connectivity patterns and role in behavioral control vary along the rostro-caudal axis of the septum^2,3,5^. The septum is populated by multiple molecularly distinct neuron subtypes, although the true extent of their diversity is still unknown^2,6,7^. To understand how the septum controls specific behavioral responses, it is necessary to study defined neuronal populations and their connectivity. Developmental lineage is often a good predictor of mature neuron identity, as neurons with shared developmental origins tend to share aspects of their circuitry and functional features^7–11^. *Nkx2.1* is a transcription factor expressed during embryonic development in forebrain progenitors within the medial ganglionic eminence / preoptic area (MGE/PoA) and the septal eminence (SE); the latter proliferative area is distinguished as part of the developing septal complex by its expression of transcription factors of the Zic family^12–14^. Septal neurons within the *Nkx2.1*-lineage are derived primarily from progenitors located in the SE, and represent 10-30% of the total neuronal population in the mature septum^10^. Cholinergic *Nkx2.1*-lineage neurons in the medial septum are essential for normal learning and memory^8^, but the role of SE-derived neuronal populations in the lateral septum has not been studied yet. In a recent study, we generated a mouse line where *Prdm16*, a gene encoding a transcriptional regulator, was deleted from cells with a developmental history of expression of *Nkx2.1*. This led to a decrease in the number of interneurons derived from the medial ganglionic eminence (MGE) in the cerebral cortex and other brain areas such as the hippocampus and the nucleus accumbens^15^. *Prdm16* is expressed exclusively in radial glial neural progenitors throughout the vast majority of the developing central nervous system^16,17^, but it is uniquely upregulated in newborn lateral septal neurons^10^. This led us to examine the conditional knockout mice, where we discovered an almost complete ablation of *Nkx2.1*-lineage neurons in the lateral, but not the medial, septum. Single-nucleus RNA sequencing experiments showed that the majority of the missing LS cells were *Crhr2*-expressing neurons; in line with this, we found that mutant mice lacked a specific set of axonal inputs to the LS positive for the neuropeptide urocortin-3 that normally form pericellular baskets onto LS *Nkx2.1*-lineage neurons. LS *Crhr2*-expressing cells drive stress-induced anxiety-like states and modulate threat responsivity^18,19^. We exposed mutant mice to a series of stress-inducing stimuli and found no evidence of general anxiety in mutant mice; rather, they exhibit increased preference for novel stimuli and exploratory behavior. Finally, we found that *Nkx2.1*-lineage neurons in the LSd are specifically activated by a stressful stimulus. Together, our data show that this population is responsive to stress, likely inhibiting exploratory drive in favor of defensive behavioral strategies when mice are under potential threat.

## RESULTS

### Ablation of *Nkx2.1*-lineage neurons in the lateral septum of *Prdm16* cKO mice

*Nkx2.1*-lineage cells constitute a significant proportion of the medial and lateral septal nuclei^7,10,11^. Based on our previous finding that newborn septal neurons express high levels of *Prdm16*^10^, we decided to examine the effect of *Prdm16* deletion within the *Nkx2.1* lineage^15^ in the septa of control (Nkx2.1Cre;Ai14, hereafter labeled as ‘WT’) and mutant (Nkx2.1Cre;Prdm16^f/f^;Ai14, labeled as ‘cKO’) mice at postnatal day 30 (P30), a young adult stage at which development is complete (**Figure 1A**). We discovered an almost complete absence of *Nkx2.1*-lineage cells (which could be identified by their expression of the fluorescent reporter tdTomato), from the lateral septum of cKO animals (**Figure 1B**). The density of tdTomato+ cells revealed a consistent loss of this population across three different positions along the rostro-caudal axis of the septum (with decreases of 92.9%, 93.8% and 83.0% in sections I, II and III, respectively; **Figures 1C,D; S1A**). This loss could be observed in all three subnuclei within the lateral septum (LSd, dorsal; LSi, intermediate; and LSv, ventral; **Figures 1B; S1B**), while the medial septum was unaffected (**Figure 1E**). In corresponding coronal sections, the area of the septum was reduced in both male and female mutant mice, particularly in posterior regions (**Figures 1F; S1D,E**). *Nkx2.1*-lineage cells normally represent 10-30% of all cells with a septal origin (as identified by immunofluorescence for Zic-family proteins) across septal nuclei^10^, Since there was no decrease in the overall density of ZIC+ cells (**Figure S1C**), this size reduction likely implies that the loss of is not compensated by any other cell lineages. *Nkx2.1*-lineage cells in the septum comprise both neurons and astrocytes^10,20,21^; to determine that the cell ablation phenotype is specific to neurons, we performed immunofluorescence staining for the astrocyte marker SOX9 (**Figure S1F**), and found no difference in astrocyte density between WT and cKO samples either in the LS or in the MS, where the majority of *Nkx2.1*-lineage astrocytes reside^7,10,20,21^ (**Figure S1G**). Collectively, these results show that we have generated a mouse model where nearly all lateral septal neurons with a history of *Nkx2.1* expression are ablated. This model could be used to explore the role of a population of lineage-defined neurons in the regulation of some of the multiple innate behaviors controlled by the lateral septum^1–3^.

**Figure 1:**
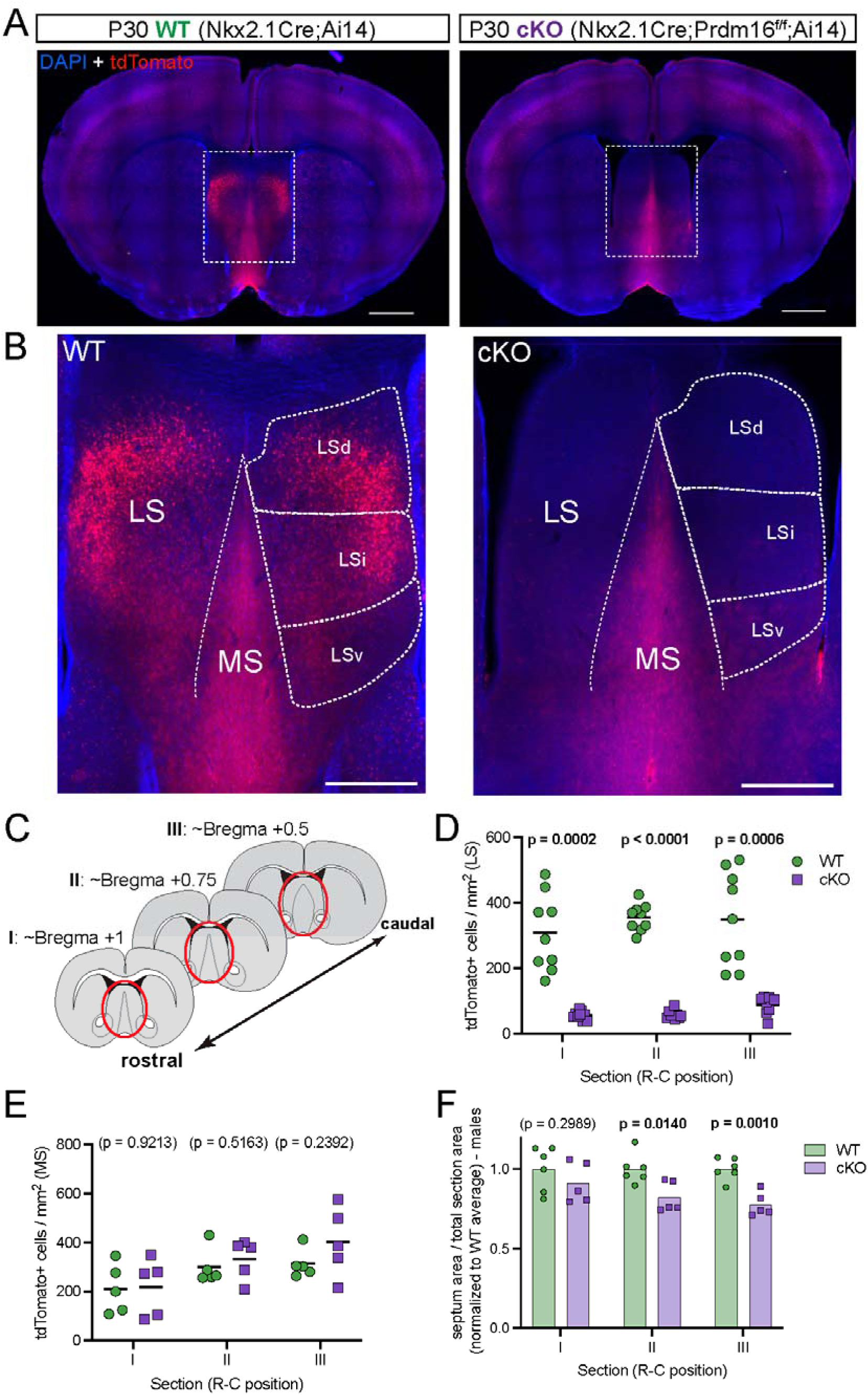
A genetic model to ablate *Nkx2.1*-lineage neurons in the lateral septum. **A)** Coronal sections through the forebrain of postnatal day 30 (P30) control (labeled as ‘WT’; Nkx2.1-Cre;Ai14, left) and mutant (labeled as ‘cKO’; Nkx2.1-Cre;Prdm16^flox/flox^;Ai14, right) mice, where cells derived from Nkx2.1-expressing progenitors are labeled by the fluorescent reporter tdTomato (red). Nuclei are counterstained with DAPI (blue). Scale bars, 1 mm. **B)** Closeup of the septum of WT (left) and cKO (right) samples, as highlighted by white dashed line boxes in **A**. The main anatomical divisions of the mature septum are indicated by white dashed lines: medial septum (MS), lateral septum (LS), and dorsal (LSd), intermediate (LSi) and ventral (LSv) nuclei within the LS. Scale bar, 500 µm. **C)** Cartoon representing forebrain coronal sections at the three rostrocaudal positions used in subsequent analyses, labeled as sections ‘I’, ‘II’ and ‘III’ throughout the article. The septal area is highlighted by red ellipses. **D)** Quantification of the density of cells positive for tdTomato per mm^2^ in the lateral septum of WT (green circles, n = 9) and cKO (purple squares, n = 9) mice. **E)** Quantification of the density of tdTomato+ cells per mm^2^ in the medial septum of WT (green circles, n = 5) and cKO (purple squares, n = 5) mice. **F)** Quantification of the area of the septum relative to the total area of its corresponding coronal brain section in WT (green circles, n = 6) and cKO (purple squares, n = 5) male mice. Measurements are normalized to the corresponding WT average. Unpaired t-tests with Welch’s correction were performed; the p-values are shown above the corresponding compared sets of data: bold typeface indicates statistically significant (p<0.05) differences.

### Loss of *Crhr2*-expressing neurons in *Prdm16* cKO mice

Recent studies have used single-cell (scRNA-Seq) and single-nucleus RNA sequencing (snRNA-Seq) techniques to describe the molecular diversity of neurons in the lateral septum, identifying multiple distinct populations based on their transcriptomic profiles, anatomical location, connectivity patterns and intrinsic electrophysiological properties^22–24^. We used this approach, and the existing knowledge derived from it, as a starting point to investigate the identity of the neurons missing from the LS of cKO mice. We generated a snRNA-Seq dataset where we dissected the septa of P35 WT and cKO mice where the Ai14 reporter allele was replaced by a different Cre-dependent reporter that expresses GFP fused to the nuclear envelope protein SUN1. This allowed us to fluorescently sort nuclei, distinguishing between the *Nkx2.1*-lineage and others (**Figure 2A**). We describe the WT samples from this dataset in detail in another study, where we found that developmental lineage is not a strict determinant of mature identity for all LS neuronal subtypes^24^. Combining 6 WT and 6 cKO samples (3 males and 3 females each, pooled to generate a single dataset), we obtained the transcriptional profiles of a total of 24323 cells (**Figure S2A**); based on their enriched expression of multiple marker genes, 8783 of those cells were classified as septal neurons (**Figure S2B**). These were then re-clustered and reanalyzed (**Figure S2C,D**), using another series of marker genes to identify a subset of 4984 lateral septal neurons, which were subdivided into multiple subtypes largely in agreement with previously published studies^22–24^ (**Figure 2B,C**). We inferred that the neuronal populations missing from the LS of cKO mice correspond to four of those neuronal subtypes, which are composed largely by *Nkx2.1*-lineage neurons (**Figure S2E**), belonging mainly (clusters 7, 8, and 9) or exclusively (cluster 13) to WT samples (**Figure 2D,E**). Expression of the genes *Esr1*, *Tshz2*, *Chat* and *Crhr2* was enriched in clusters 7, 8, 9, and 9 + 13, respectively (**Figure 2F,G**), especially in *Nkx2.1*-lineage nuclei (**Figure S2F**). We focused on *Crhr2*, a gene encoding the type 2 corticotropin-releasing hormone receptor, as the LS contains a neuronal population expressing this gene that plays a crucial role in anxiety-like behavior and threat avoidance in rodents^18,19,25^. We performed *in situ* hybridization for *Crhr2* in WT and cKO mice at P30 (**Figure 2H,J**), and found a decrease in the number of *Crhr2*-expressing cells in the LS of cKO mice, which was due to the almost complete ablation of the *Nkx2.1*-lineage within this population (**Figures 2J, S2G**). The majority of LS *Crhr2*-expressing cells are confined to the tdTomato+ *Nkx2.1*-lineage in WT brains (**Figure S2H**), indicating that their developmental origin is the septal eminence^10^. Detailed quantifications show that the density of *Crhr2*-expressing cells in the LS of WT samples is highest in middle rostro-caudal positions (i.e. section II), increasing in a dorsal to ventral pattern (**Figure S2I**). *Crhr2*+ cell density is unaffected in the caudal sections of cKO brains (**Figure S2J**), where the proportion of *Nkx2.1*-lineage cells within this population is lowest (**Figure S2K**). The increase in the density of *Crhr2*+, tdTomato–cells in cKO animals suggests that there might be a partial compensatory effect within this population, whereby cells from different developmental origins would be able to respond to the absence of *Nkx2.1*-lineage cells by turning on *Crhr2* expression. However, it is unclear whether this apparent change in gene expression from other neuronal populations could restore any functional defects brought by the loss of SE-derived LS neurons. We began to address this question by analyzing the connectivity of the LS in WT and cKO animals.

**Figure 2:**
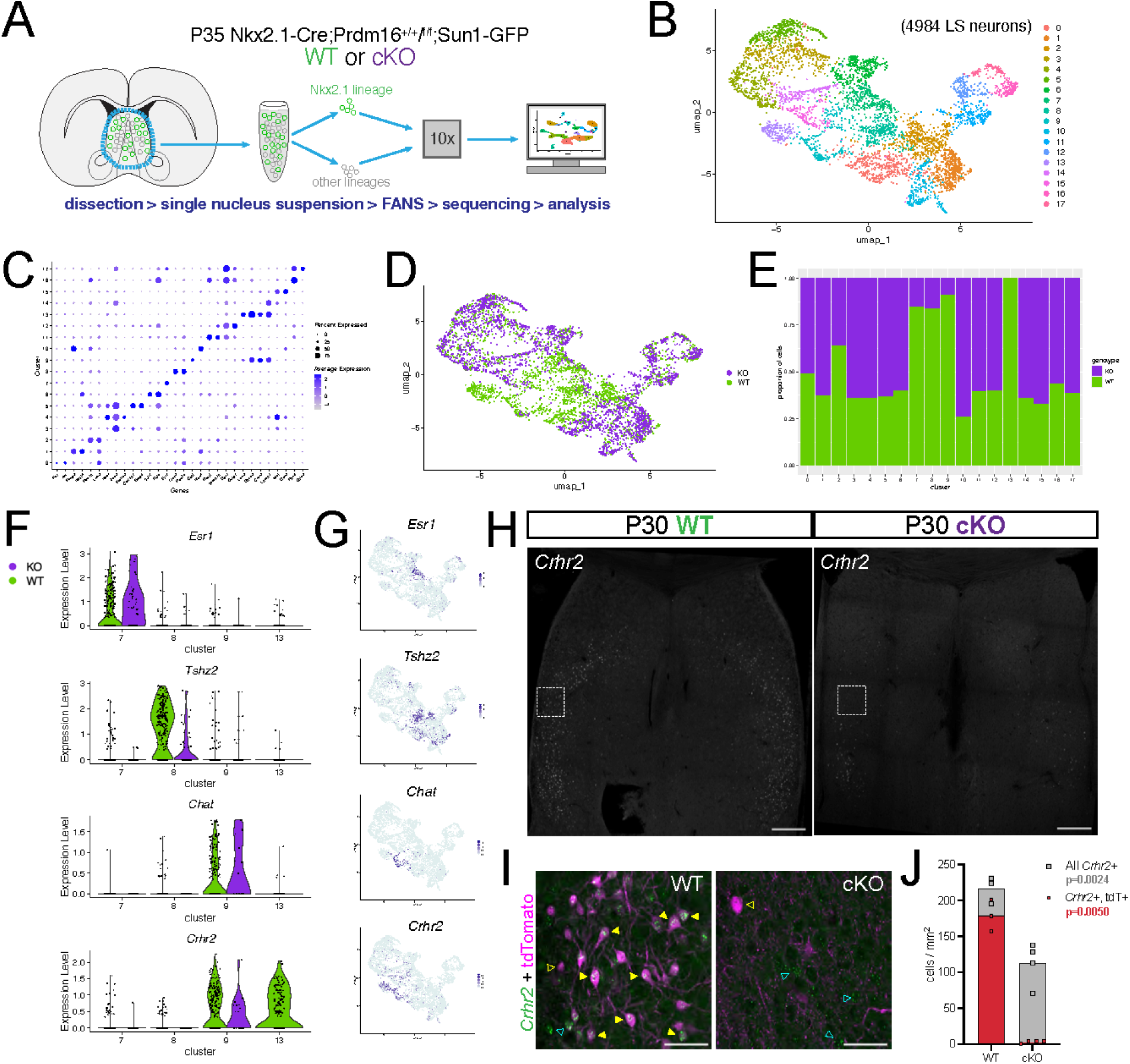
Loss of *Crhr2*-expressing neurons in cKO mice. **A)** Cartoon illustrating the experimental strategy. The septa of P35 ‘WT’ (Nkx2.1-Cre;Sun1-GFP) or ‘cKO’ (Nkx2.1-Cre;Prdm16^flox/flox^;Sun1-GFP) animals carrying a Cre-dependent fluorescent reporter inserted into the nuclear membrane were dissected out, sorted into Nkx2.1+ vs. Nkx2.1– populations via Fluorescence-Activated Nucleus Sorting (FANS), and submitted to single-nucleus RNA sequencing using the 10x platform. **B)** Uniform Manifold Approximation and Projection (UMAP) plot representing a 2D dimensional reduction of the transcriptional similarity among all cells identified as LS neurons within the dataset (4984 cells from a total of 6 WT and 6 cKO animals [3 males and 3 females each]). **C)** Dot plot showing the expression of select marker genes in each of the LS clusters outlined in **B**, allowing their identification as distinct cell types. **D)** UMAP plot as in **B**, with sample identities highlighted as indicated (WT, green; cKO, purple). **E)** Bar graph showing the proportion of cells belonging to WT (green) *vs*. cKO (purple) samples within each of the 17 clusters outlined in **B**. **F)** Violin plots showing the expression levels of select marker genes within clusters 7, 8, 9, and 13 (i.e. all clusters where ≥75% of cells belong to WT samples), split by genotype (WT, green; cKO, purple). **G)** Feature plots showing the gene expression levels within the UMAP representation. **H)** Overview coronal images of the septum of P30 WT (left) and cKO (right) mice submitted to *in situ* hybridization for *Crhr2* (gray). Scale bars, 250 µm. **I)** Closeup view of the white dashed line boxes in **H**, showing the *Crhr2* signal (green) combined with immunofluorescence staining for tdTomato (magenta), in the LS of WT (top) and cKO (bottom) mice. Full yellow arrowheads indicate examples of cells positive for both *Crhr2* and tdTomato, empty yellow arrowheads show examples of *Crhr2*–, tdTomato+ cells, and empty cyan arrowheads highlight examples of *Crhr2*+, tdTomato– cells. Scale bars, 50 µm. **J)** Bar graph showing the density (per mm^2^) of all cells positive for *Crhr2* (gray squares) and the subset of those cells that are also tdTomato+ (red circles), in section II of WT *vs*. cKO animals (N = 3 [2 males, 1 female] for WT; N = 4 [2 males, 2 females] for cKO).

### Reduced innervation of the lateral septum in the absence of *Crhr2*-expressing neurons

The lateral septum is densely innervated by axons positive for urocortin-3 (UCN-3), a neuropeptide within the corticotropin-releasing factor family with high affinity for the CRHR2 receptor, whose activation has an anxiogenic behavioral effect^1,26–28^. We reasoned that the absence of *Crhr2*-expressing neurons in the LS of cKO mice could result in circuit disruptions derived from an altered distribution of axonal terminals containing UCN-3. We performed immunofluorescence staining for UCN-3, and found that while UCN-3 terminals densely innervated the LS of WT brains, they were largely absent in cKO mice (**Figure 3A,B**). We quantified the fluorescence intensity of UCN-3 staining, confirming that the reduced innervation in cKO mice was consistent across the rostro-caudal and dorso-ventral axes, except for the LSv of the most rostral section (**Figure 3C**). UCN-3 axonal terminals form perineuronal baskets around the somata of tdTomato+ neurons in control conditions, indicating that these inputs are specific to *Nkx2.1*-lineage cells. This was not the case in cKO samples, where the few remaining UCN-3+ baskets were largely not specific to tdTomato+ cells (**Figure 3B,D-F**). We validated the observed decrease in innervation using Enkephalin (Enk), another neuropeptide that highly overlaps with UCN-3 in axonal terminals throughout the LS^6,29–31^. We compared immunofluorescence staining for Enk in WT and cKO brains (**Figure S3A,B**), finding a decrease in cKO samples that paralleled our results for UCN-3 (**Figure S3C,D**). To confirm that this disruption in connectivity was specific to UCN-3/Enk inputs onto *Nkx2.1*-lineage cells, we performed immunofluorescence staining for other known inputs to the LS, including tyrosine hydroxylase (TH), as a proxy for dopamine^32^ (**Figure S3E**), and serotonin (5-HT; **Figure S3F**)^33,34^, and did not find any abnormal patterns in cKO animals. These concurrent observations provide compelling evidence that the connectivity of the lateral septum of cKO mice is disrupted specifically due to the absence of the *Crhr2*-expressing *Nkx2.1*-lineage neuronal population normally targeted by UCN-3/Enk inputs.

**Figure 3:**
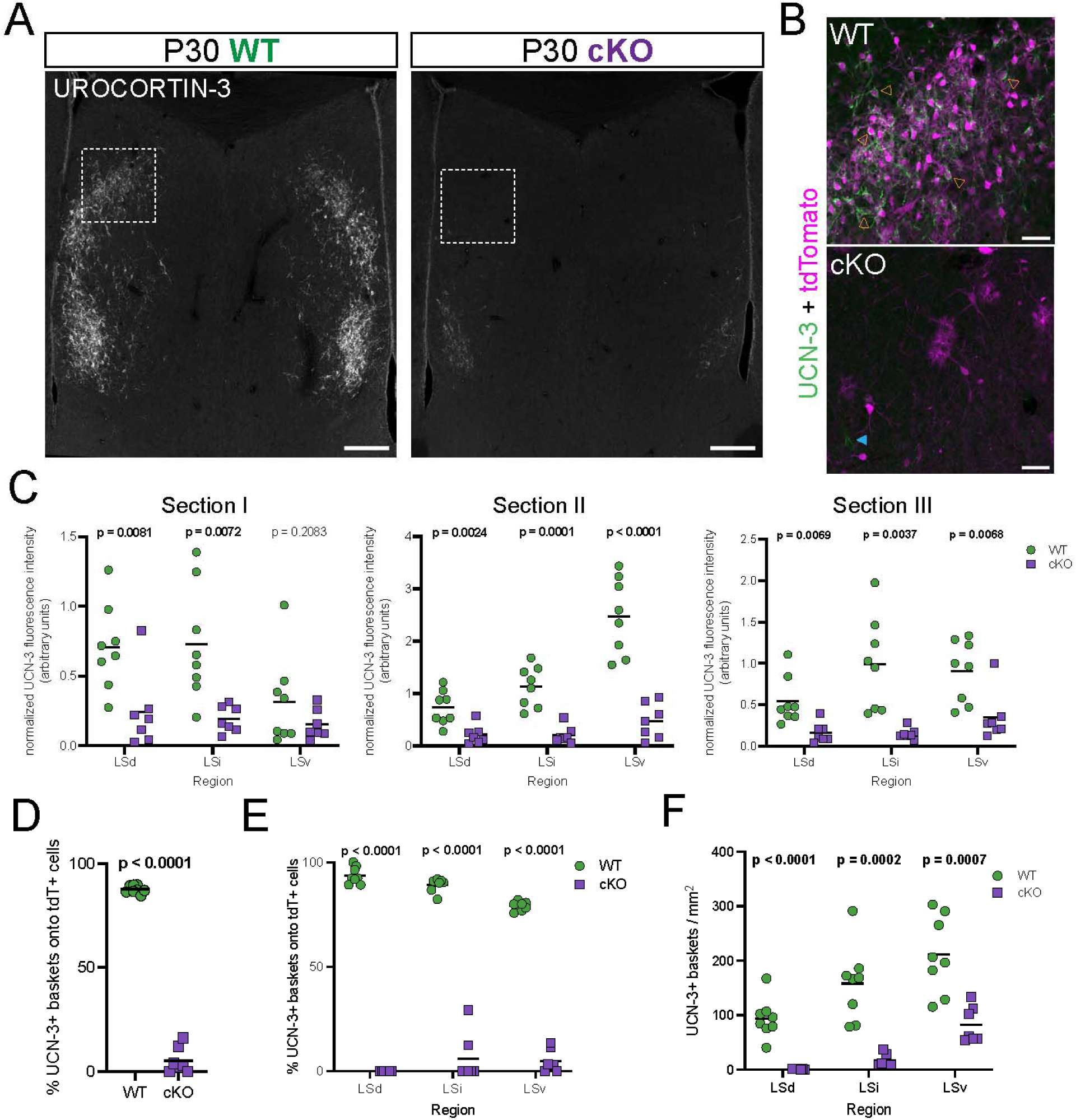
Disrupted connectivity in the LS of cKO mice. **A)** Overview coronal images of the septum of P30 WT (left) and cKO (right) mice submitted to immunofluorescence staining for urocortin-3 (gray). Scale bars, 250 µm. **B)** Closeup view of the white dashed line boxes in **A**, showing the urocortin-3 signal (UCN-3, green) combined with tdTomato (magenta), in the LS of WT (top) and cKO (bottom) mice. Empty orange arrowheads indicate examples of tdTomato+ cells surrounded by urocortin-3 perineuronal baskets, while the blue arrowhead shows a basket formed around a tdTomato– cell. Scale bars, 50 µm. **C)** Quantification of normalized fluorescence intensity in the different LS subnuclei of P30 WT (green circles) and cKO (purple squares) samples where urocortin-3 was detected by immunofluorescence staining. **D)** Quantification of the proportion (in %) of all urocortin-3 perineuronal baskets surrounding tdTomato+ cells throughout the entire LS of WT (green circles) and cKO (purple squares) mice at P30. **E)** Breakdown of the data in **D** by LS subnucleus. **F)** Quantification of the density of urocortin-3 baskets per mm^2^ in the different LS subnuclei of P30 WT (green circles) and cKO (purple squares) mice. **C-F:** N = 8 (4 males, 4 females) for WT; N = 7 (3 males, 4 females) for cKO. Unpaired t-tests with Welch’s correction were performed; the p-values are shown above the corresponding compared sets of data: bold typeface indicates statistically significant (p<0.05) differences. **E**, **F** and **G** show quantifications in rostro-caudal section II, as defined in **Figure 1C**.

### Mild alterations in the electrophysiological properties of LS *Nkx2.1*-lineage neurons in cKO mice

Next, we sought to understand how the loss of an entire population of neurons in the LS and its associated disruptions in connectivity might affect neuronal function in cKO mice. We started by examining the intrinsic electrophysiological properties of LS neurons, performing whole-cell patch clamp recordings in cells within the *Nkx2.1*-lineage (tdTomato+) and outside of it (tdTomato–), in both WT and cKO samples (**Figure 4A**). Most intrinsic membrane properties, including resting membrane potential, input resistance, spike half-width and threshold, adaptation ratio, afterhyperpolarization and rheobase (**Figures 4D,E; S4A,E-G**), were similar for neurons of both lineages across genotypes. When we analyzed the firing patterns and action potential waveforms (**Figure 4B,C**), we observed that *Nkx2.1*-lineage neurons fired smaller initial spikes compared to neurons outside of this lineage. The time it took to fire those initial spikes was extremely variable in *Nkx2.1*-lineage neurons in cKO samples, showing a greater average latency compared to their WT counterparts and to neurons outside of this lineage (**Figure S4C,D**). Membrane capacitance was lower in in cKO samples (**Figure 4F**), but only correlated with a lower membrane time constant in non-*Nkx2.1*-lineage cells (**Figure S4B**). The afterhyperpolarization of *Nkx2.1*-lineage neurons was statistically greater in cKO cells, reflecting their higher variability (**Figure S4H**). When we subjected patched cells to step-wise increasing current injections, we observed that while the tdTomato– population responded similarly across genotypes, *Nkx2.1*-lineage neurons in cKO samples fired action potentials at higher frequencies; this difference was statistically significant for currents of 100 pA and above (**Figure 4B,C,G**), which indicates that the remaining *Nkx2.1*-lineage neurons are more excitable in cKO mice. These observations show that only a few of the intrinsic electrophysiological properties of *Nkx2.1*-lineage neurons in the LS are distinct from those of the surrounding populations. This appears especially true in cKO samples, whose greater variability, lower membrane capacitance, and increased firing rate at high current injections might reflect intrinsic molecular differences (such as different channel composition) and/or circuit defects (e.g. decreased or deficient inhibitory tone due to the loss of GABAergic neurons that might normally form local LS circuits within the *Nkx2.1*-lineage). Most of the neurons that we observed, both in WT and cKO samples, would be classified as “tonic cells” according to previous studies^35–37^; however, it is possible future experiments could reveal a further diversity of firing patterns, reflecting potential different electrophysiological subtypes of neurons within the *Nkx2.1*-lineage as we have observed before^24^.

**Figure 4:**
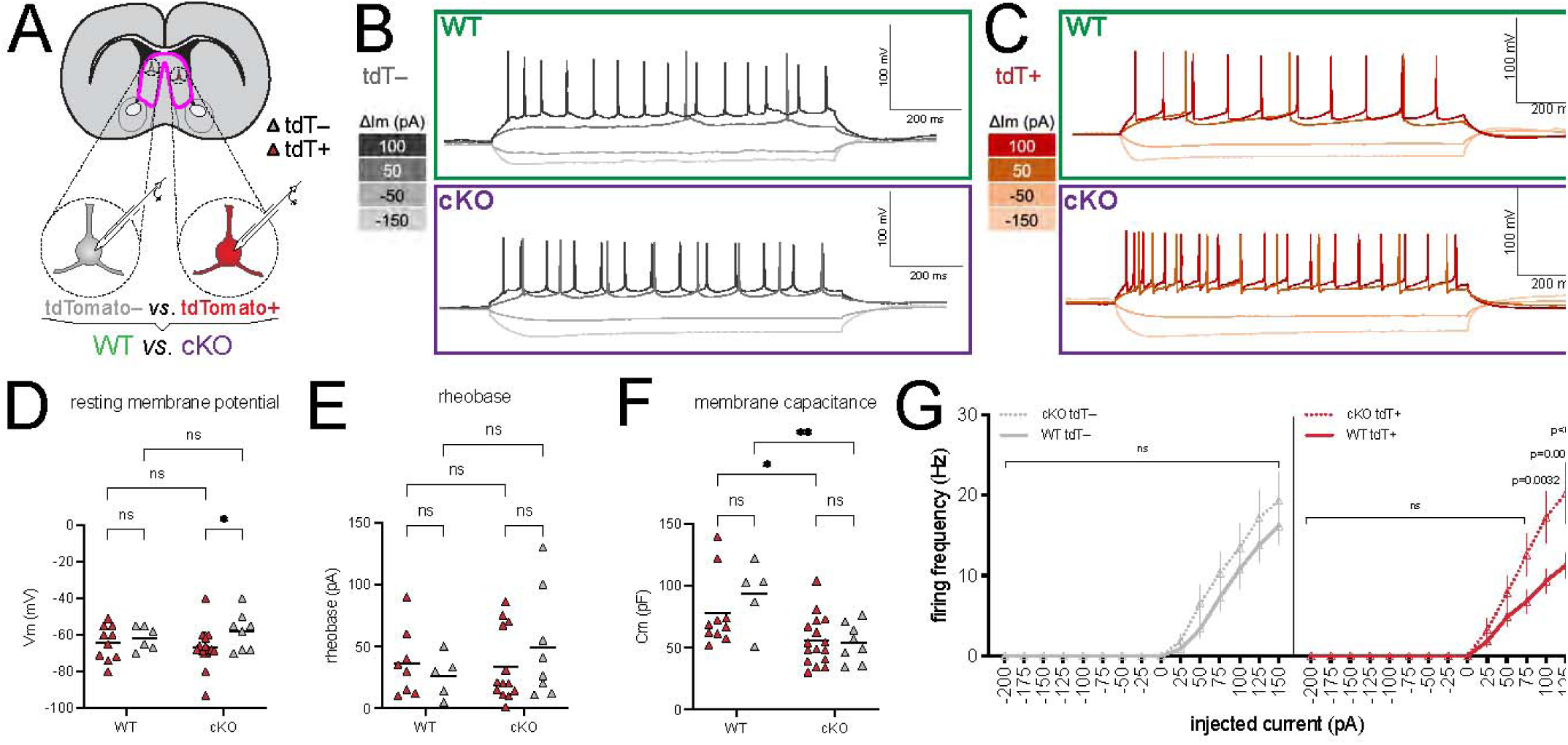
Intrinsic electrophysiological properties of WT and cKO LS neurons. **A)** Schematic of the experimental setup: whole-cell recordings were obtained from lateral septal neurons, both tdTomato+ and tdTomato–, on slices from WT and cKO animals at 3 weeks of age. **B, C)** Representative traces of the firing patterns of tdTomato– (**B**) and tdTomato+ (**C**) neurons of WT (top) and cKO (bottom) samples subjected to current injection at the indicated intensities. **D-F)** Comparison of intrinsic electrophysiological properties of LS neurons of WT and cKO samples: **D**, resting membrane potential; **E**, membrane capacitance; **F**, rheobase. **G)** Comparison of the firing frequency of tdTomato– (left) and tdTomato+ (right) LS neurons subjected to step-wise current injections at 25 pA intervals, from -200 to 150 pA (data are represented as average +/-SEM). For WT tdTomato+, n = 10 (D, E), n = 9 (F), n = 16 (G); for WT tdTomato–, n = 6 (D), n = 5 (E, F), n = 8 (G); for cKO tdTomato+, n = 16 (D, E), n = 13 (F), n = 7 (G); for cKO tdTomato–, n = 8 (D-F), n = 10 (G). After removal of outliers via Grubbs’ test, 2-way ANOVA tests with multiple comparisons were performed; the p-values are shown above the corresponding compared sets of data: bold typeface indicates statistically significant (p<0.05) differences.

### Increased exploratory drive in cKO mice

We decided to investigate the possible consequences of the loss of LS *Nkx2.1*-lineage neurons at the systemic level. Previous research has shown that *Crhr2*-expressing LS neurons modulate behavioral responses to threatening stimuli when activated, producing an anxiety-like state when mice are exposed to persistent stress^18,19^. We subjected control and mutant animals to a series of behavioral tests designed to assess general anxiety-like behaviors in mice that are mediated at least partially by the LS^18,38^ (**Figure S5A**). Our results did not indicate a heightened or decreased anxiety status in cKO mice, as shown by classic paradigms such as the open field test (**Figure 5A**) or the elevated plus-maze test (**Figure S5B**), or any defects in social interaction (**Figure S5C**) or spatial memory (**Figure S5D**). We only observed significant differences between genotypes in the novel object and light-dark box paradigms: male cKO mice spent more time exploring the novel object in the former (**Figure 5B**), and more time in the dark chamber in the latter (**Figure 5C**). We reasoned that the congenital ablation of *Nkx2.1*-lineage LS neurons might result in partial behavioral compensation by other circuits, such that observing clear differences between WT and cKO mice might require a more salient and aversive stimulus. We chose to perform a predator odor test, exposing mice to the smell of TMT (2,3,5-trimethyl-3-thiazoline), a component of fox odor that triggers an innate and strongly aversive response mediated by neurons in the lateral septum that are highly likely to belong to the *Nkx2.1*-lineage based on their anatomical distribution^39,40^. To carry out the test, mice were placed in an arena equipped with an air inlet, and subjected to two experimental phases: an initial 15-minute habituation phase (‘Blank phase’), similar to an open-field test, in which the arena was filled with air carrying a neutral odor. This was followed by a subsequent 15-minute test phase, during which the air in the arena contained the smell of TMT (‘TMT phase’)^41,42^ (**Figure 5D**). The behavior of mice throughout both phases was recorded with a depth sensor camera located above the center of the arena. Mice of both genotypes and sexes spent time exploring the novel stimulus presented by the air flow from the inlet during the Blank phase, while they tended to move away from it during the TMT phase, as shown in the corresponding occupancy plots (**Figure 5E**). cKO animals displayed a higher baseline exploratory drive, as they spent a higher proportion of time around the odor inlet in the Blank phase of the experiment. This proportion was reduced in all animals during the TMT phase (i.e., after they were exposed to the predator smell), but cKO mice continued to explore the air inlet for longer than their WT counterparts, especially in the case of male animals (**Figure 5E,F**). This suggests that, within this behavioral paradigm, novelty-triggered exploration can override anxiety-induced aversion in mutant mice. To further analyze behavioral differences between groups, we used MoSeq, an unsupervised automated behavior analysis pipeline^41^. This method detects sub-second behavioral motifs (or ‘syllables’) and quantifies their usage, facilitating unbiased comparisons across conditions to guide detailed analyses of behavior^41–43^. We observed certain emerging patterns when comparing the usage of each syllable in Blank *vs*. TMT phases within each genotype and sex. Syllables associated with ambulatory and/or exploratory behaviors (such as darting or rearing) were comparatively enriched during the Blank phase, while those related to stationary behavior (such as pausing or grooming) were used more often during the TMT phase (**Figure S5E,F,I,J**). This was consistent for both sexes and genotypes, and reflected our observations outlined above regarding the typical behavior that mice displayed during this experiment. We also compared syllable usage between WT and cKO mice in both the Blank and TMT phases, to assess the behavioral repertoire of each genotype in the absence and presence of a threatening stimulus. During the Blank phase, female WT mice displayed heightened vigilance, with increased frequencies of behavioral motifs such as jumping (syllable 26) and quick rearing (syllable 33). Conversely, female cKO animals differentially exhibited behaviors such as post-rearing ambulation (syllable 13) and prolonged, high rears (syllable 25), reflective of exploration (**Figure S5G**). In the TMT phase, animals of either genotype tended to use similar syllables differentially, suggesting that the heightened state of alert *vs*. exploration is consistent when comparing WT and cKO mice, respectively (**Figure S5H**). Male mice, despite their similarly strong response to TMT (**Figure 5E,F**), showed more subtle differences in syllable usage between WT and cKO animals (**Figure S5K,L**). Taken together, the results from these experiments show that, while animals of both genotypes exhibit threat avoidance behavior, cKO mice display an increased exploratory drive even when faced with a salient stressor.

**Figure 5:**
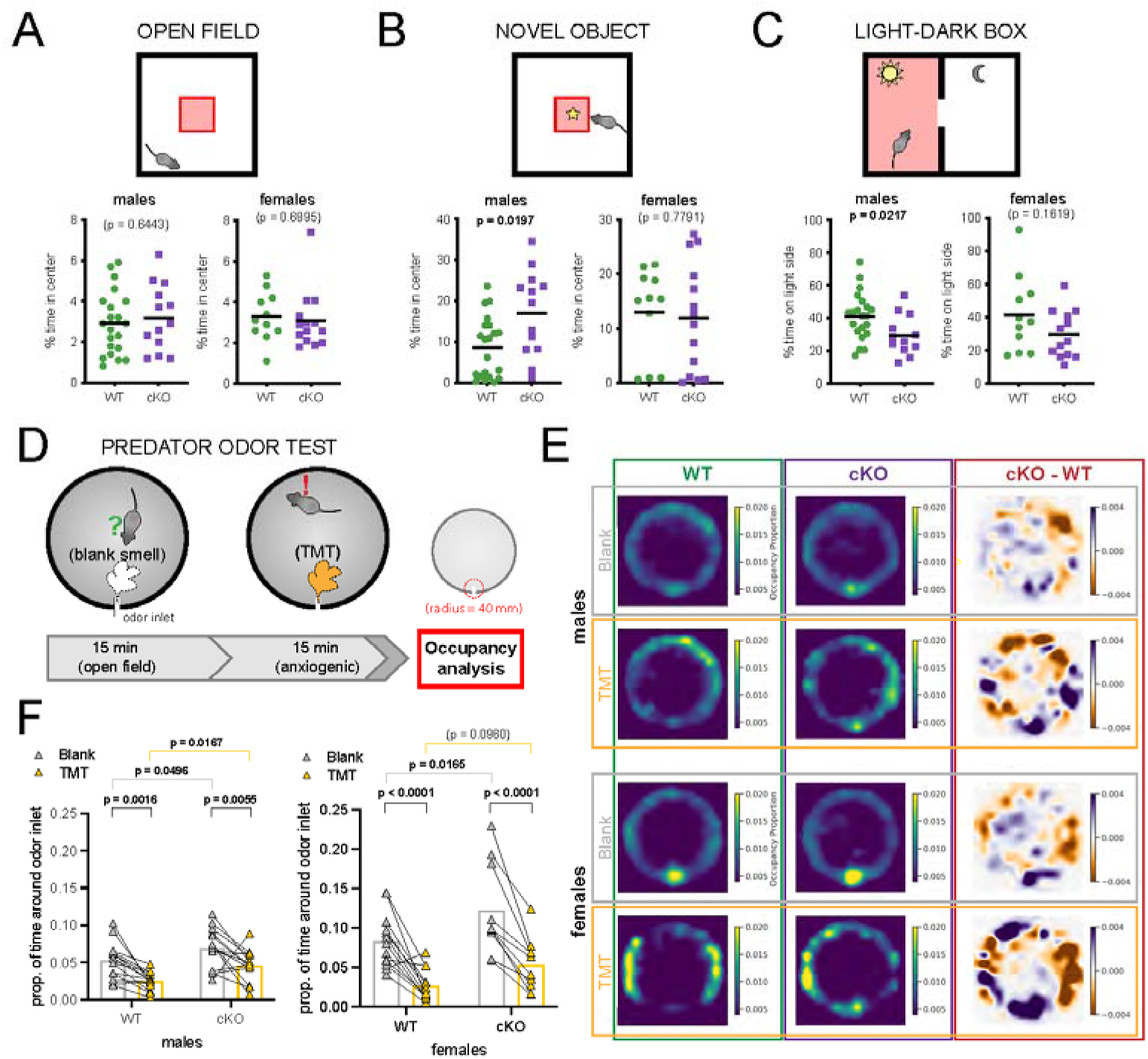
cKO mice display increased exploratory drive. **A-C)** Top, cartoons illustrating the outline of each behavior test; bottom, summary of the main anxiety-related readout for both males and females. The behavioral tests performed were: **A**, open field; **B**, novel object; **C**, light-dark box. Unpaired t-tests with Welch’s correction were performed; the p-values are shown above the corresponding compared sets of data: bold typeface indicates statistically significant (p<0.05) differences. **D)** Experimental design: mice were placed in a cylindrical arena with a single odor inlet near the bottom, and exposed to two consecutive phases, where either a neutral (Blank) or an aversive (TMT) smell was pumped into the arena. **E)** Occupancy plots showing the average proportion of experiment time spent in each location throughout the arena by WT (n = 15 males, 13 females) or cKO (n = 14 males, 10 females) mice, under blank odor (“Blank”) or anxiogenic (“TMT”) conditions, as well as the difference between both plots (“cKO-WT”), separated by sex (top, males; bottom, females). **F)** Plot displaying the average occupancy of the area within a 40 mm radius around the odor inlet during Blank (gray triangles) and TMT (golden triangles) conditions, separated by sex and genotype. Each line joins the occupancy values of an individual mouse. Bars indicate the average. Note the same range in the y-axis of males and females. After removal of outliers via Grubbs’ test, 2-way ANOVA tests with multiple comparisons were performed; the p-values are shown above the corresponding compared sets of data: bold typeface indicates statistically significant (p<0.05) differences.

### *Nkx2.1*-lineage neurons in the LS are activated by acute restraint stress

Given the strong connection between LS *Crhr2*-expressing neurons and anxiety-like behavior^18,19^, and between predator odor response and the lateral septum^39,40,44^, we reasoned that the behavioral differences between WT and cKO mice that we observed were likely to be a consequence of the loss of LS *Nkx2.1*-lineage neurons in mutant mice. However, the experiments described above did not test if this population was specifically activated when mice were faced with a threatening stimulus. To test if this was the case, we subjected WT and cKO mice to acute forced restraint, a highly aversive stimulus that induces a strong activation of LS neurons^45–47^. Mice were restrained for 30 minutes and sacrificed one hour later to analyze the activation of LS cells, using immunofluorescence staining for c-Fos as a readout (**Figure 6A**). The lateral septum of animals submitted to forced restraint displayed numerous c-Fos+ neurons (identified by the neuronal marker NeuN), in both WT and cKO samples (**Figure 6B,D**). We quantified the density of c-Fos+ neurons throughout the LS along the dorsolateral axis, and found a robust increase in experimental animals, compared to the baseline in untreated mice (**Figures 6D, S6A,B**). As expected, the proportion of c-Fos+ cells belonging to the *Nkx2.1* lineage (i.e., tdTomato+) was consistently higher in WT than cKO brains, in all LS subnuclei (**Figures 6C, S6C**). The proportion of *Nkx2.1*-lineage cells within the c-Fos+ population across the LS was significantly higher in animals subjected to forced restraint than in untreated controls (**Figure 6E**); this was true even in cKO animals, where the overall proportion of tdTomato+ cells was much lower (**Figure S6D**). This increased proportion indicates that anxiogenic stimuli specifically activate *Nkx2.1*-lineage neurons in the lateral septum, consistent with the pattern of c-Fos staining in previous studies where animals were subjected to various anxiety-inducing paradigms^39,48^. As a result of the ablation of this lineage, we observed a statistically significant reduction in the density of c-Fos+ neurons specifically in the LSd (**Figure 6F**), but not in the LSi or LSv, of cKO mice compared to WT (**Figure S6E**). These results demonstrate that a highly aversive stimulus activates LS *Nkx2.1*-lineage neurons, pointing to a role for this population in the response to anxiogenic environments that is consistent with the behavioral observations outlined above.

**Figure 6:**
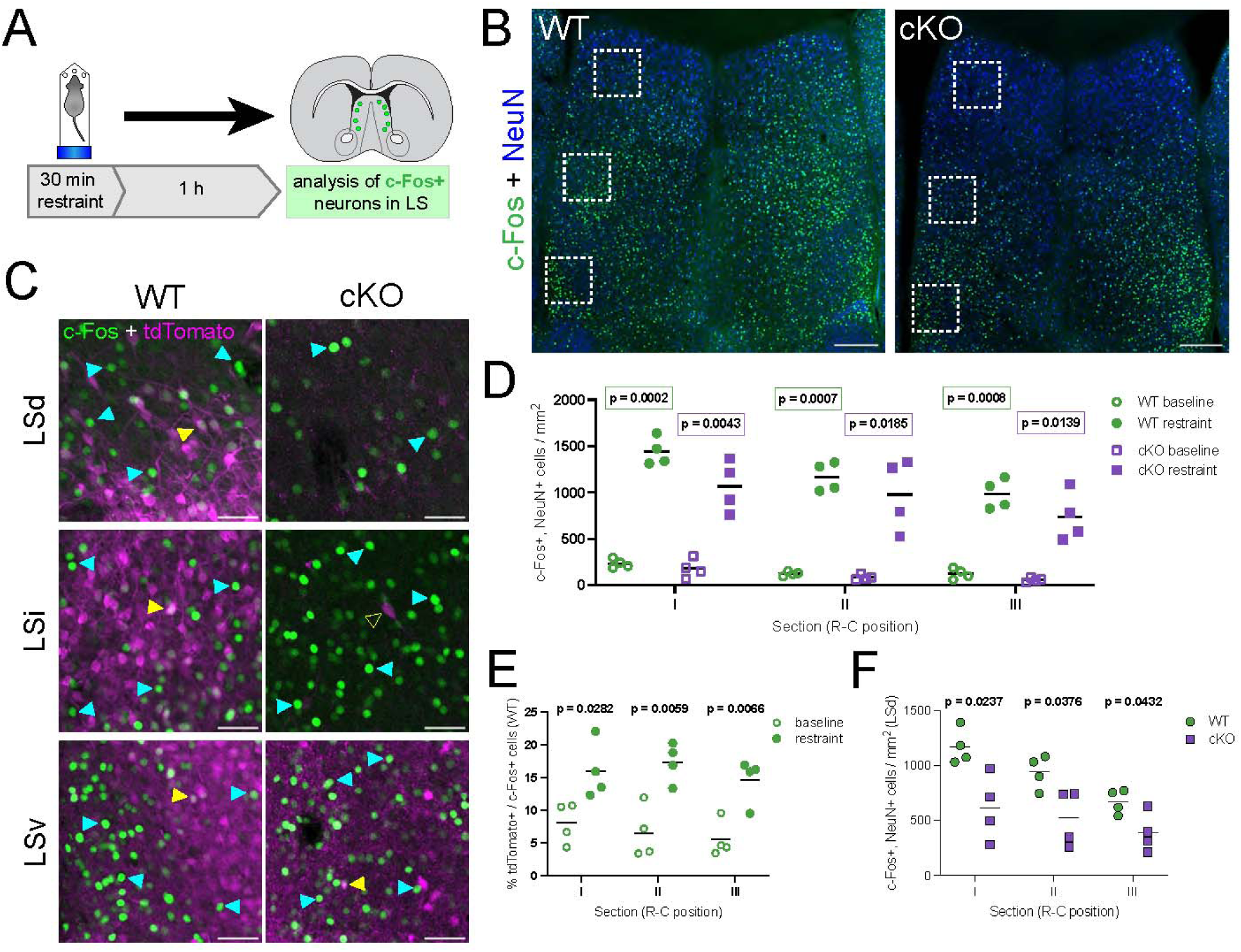
N*k*x2*.1*-lineage neurons in the LSd are activated by an acute stressful stimulus. **A)** Experimental design: mice were subjected to 30 minutes of forced restraint and sacrificed 1 hour later; neurons firing in response to the anxiogenic stimulus were identified by immunofluorescence staining for c-Fos. Scale bars, 250 µm. B) Examples of coronal sections of the septum of WT (left) and cKO (right) mice after the forced restraint experiment, immunostained for c-Fos (green) and NeuN (blue). **C)** Closeup view of the white dashed line boxes in **B**, showing c-Fos (green) and tdTomato (magenta), in the different subnuclei within the LS of WT (left) and cKO (right) mice. Yellow arrowheads indicate examples of cells positive for both c-Fos and tdTomato; cyan arrowheads highlight c-Fos+, tdTomato– cells, and the empty yellow arrowhead shows an example of a c-Fos–, tdTomato+ cell. Scale bars, 50 µm. **D)** Comparison of the density of c-Fos+ neurons in the LS of untreated controls (‘baseline’, empty symbols; n = 4 WT, green circles; n = 4 cKO, purple squares) and animals subjected to forced restraint (‘restraint’, full symbols; n = 4 WT, green circles; n = 4 cKO, purple squares). **E)** Proportion of tdTomato+ neurons within the c-Fos+ population in the LS of WT mice, comparing untreated controls (‘baseline’, empty circles, n = 4) and animals subjected to forced restraint (‘restraint’, full circles, n = 4). **F)** Comparison of the density of c-Fos+ neurons in the LSd of WT (green circles, n = 4) and cKO (purple squares, n = 4) animals subjected to forced restraint. Unpaired t-tests with Welch’s correction were performed; the p-values are shown above the corresponding compared sets of data: bold typeface indicates statistically significant (p<0.05) differences.

## DISCUSSION

The lateral septum plays a central role in the regulation of numerous emotional and motivational behavioral states^1–3^. However, attributing particular aspects of behavioral control to distinct LS neuronal subpopulations has been challenging due to the incomplete understanding of LS neuronal diversity. Recent studies have focused on several molecularly defined groups of LS neurons to establish their involvement in social behavior^49,50^ or context-specific defensive responses^18,19,51,52^. Our research complements these efforts, using a mouse model where *Crhr2*-expressing *Nkx2.1*-lineage neurons are absent from the lateral septum to assess the role of this subpopulation in the control of innate behavioral responses to stress-related stimuli. In line with previous research in the MS, we show that developmental origin is a strong predictor of septal neuron identity, and that lineage-based ablation of defined neuronal populations provides a valuable tool to test their function in the mature septum^8,11^.

Deletion of *Prdm16* in *Nkx2.1*-expressing progenitors leads to the loss of approximately 30% of cortical interneurons derived from the MGE/PoA, whose function is partially compensated by other interneuron types^15^. This likely the result of reduced levels of apoptosis in cortical interneurons derived from different developmental lineages, as suggested by observations in similar models^53^. By contrast, septal eminence-derived *Nkx2.1*-lineage neurons in the LS appear to be especially susceptible to the loss of *Prdm16*, since it results in their almost complete ablation (**Figures 1, S1**). This is accompanied by a decrease in the size of the septum, suggesting that other progenitor lineages in the developing cKO brain are unable to increase their neuronal output or survival to compensate for the loss of LS *Nkx2.1*-lineage neurons. However, it is possible that other neuron types might be rewired in cKO brains, participating in new circuits and potentially exerting some degree of functional compensation for the missing the *Nkx2.1* population. Future research establishing direct comparisons between both genotypes and refining the existing connectivity maps of the LS will be necessary to address this and gain a better understanding of the results of the behavioral tests we conducted.

In the MGE/PoA, expression of *Prdm16* is confined to progenitors, where it controls their proliferative capacity^15^. In the developing septum, *Prdm16* expression is sustained in postmitotic neurons^10,54^, potentially reflecting a specific need for this gene in the survival of septal neurons. The *Nkx2.1*-lineage in the LS is largely derived from the septal eminence, which produces sequential cohorts of neurons with diverse morphologies and anatomical locations over the neurogenic period^10^. This suggests that this lineage is composed of various neuron subtypes with distinct connectivity patterns and mature functions^2^. Besides their common developmental origin, mature *Nkx2.1*-lineage neurons share widespread expression of *Crhr2*, which in turn is largely confined to this population (**Figures 2, S2**). The functional organization of cortical circuits depends heavily on the generation of diverse cell types that retain certain shared features based on their developmental origins^55^. Like the cortex, the lateral septum is composed of diverse cell types derived from multiple progenitor domains^7,10,11^. The confluence of different developmental lineages within any given brain area is likely to be an evolutionary mechanism to refine and/or expand its circuitry and behavioral control. In the context of the *Nkx2.1*-lineage in the LS, future research should address the gene regulatory networks guiding the specification of neurons derived from the septal eminence, with special focus on the potential interplay between *Nkx2.1* and *Prdm16*.

The disruptions in septal connectivity that we observed in cKO animals (**Figures 3, S3**) show that urocortin-3 inputs to the LS are specifically affected by the loss of their high affinity receptor CRHR2^26,27^, which is normally expressed by *Nkx2.1*-lineage neurons. In an earlier version of this manuscript, we reported an anecdotal observation that we have since removed due to low experimental numbers: using non-specific retrograde labeling of the LS, we identified the perifornical area of the hypothalamus (PeFAH) as the potential source of the disrupted UCN-3 inputs to *Nkx2.1*-lineage neurons in the LS. This is particularly compelling, as this area is bidirectionally connected to the septum^56–58^, and PeFAH UCN-3 neurons are activated when mice engage in novelty exploration or are exposed to acute restraint stress^56,59^, while a neighboring GABAergic neuron population in the anterior hypothalamic nucleus (AHN) that receives *Crhr2*-expressing LS inputs^18,19^ is activated during behavioral avoidance in anxiogenic contexts, including novel object approach and exposure to fox urine smell^60^. More thorough circuit tracing experiments would be necessary to explore the possibility that *Nkx2.1*-lineage neurons are part of a PeFAH>LS>AHN circuit that is responsible for risk assessment under stressful circumstances. In this scenario, the absence of PeFAH inputs to the LS in cKO animals would result in an impairment of long-range LS>AHN inhibition, either directly or through intraseptal inhibitory sub-circuitry^2^, leading to a release of behavioral avoidance and increased exploratory drive.

The lateral septum is regarded as a center that calibrates the intensity of behavioral responses to external stimuli, depending on context and on how and which of the complex intraseptal circuits are recruited^2,3^. The mouse model we describe here presents an opportunity to further understand how a subgroup of LS neurons regulates specific aspects of behavior. Based on a previous study that described the activity of *Crhr2*-expressing neurons as anxiogenic^18^, we hypothesized that cKO mice, as they lack this population^18^, might display lower levels of anxiety-related behaviors. However, we did not find clear differences in general anxiety levels between WT and cKO mice using classic behavioral paradigms (**Figure S5**), possibly due to the lack of a persistent stressor. The only significant results in these tests appear contradictory: more time spent on the dark chamber in the light-dark box test typically denotes heightened anxiety, while more time exploring the novel object in the corresponding test is regarded as an indicator of reduced anxiety. However, these observations can be reconciled if we consider that mice were placed inside the light compartment of the light-dark box test during their daytime hours (see **Methods**). Both observed behaviors can be interpreted as an increase in the exploratory drive of cKO animals when faced with a novel stimulus, preferring the unfamiliar dark environment during daytime and showing a pronounced interest in a novel physical object within a familiar setting. The observed lack of preference for spatial novelty, as seen in the Y-maze test, could reflect that this increase in exploratory behavior is dependent on the saliency of the stimulus. Using a different experimental paradigm that presents mice with the smell of a predator, a strongly aversive and anxiogenic stimulus, we observed an increased exploratory drive in cKO animals, reflected both in the greater amount of time they spent around the odor inlet and in their higher usage of exploration-related behavioral ‘syllables’ as revealed by MoSeq (**Figures 5, S5**). The lateral septum has been implicated in novelty-seeking behaviors in the past, mainly in the context of social novelty and in relation to its connectivity to other brain areas, such as the amygdala, prefrontal cortex and hypothalamus^61–63^. Our mouse model suggests that LS *Nkx2.1*-lineage neurons and the circuits they participate in are necessary to maintain the balance between defensive/avoidance behavior and exploratory drive, in line with the proven role of several other LS neuronal subpopulations and their circuitry in context-dependent risk assessment^18,19,51,52,64,65^.

Another important consideration is the loss of other forebrain *Nkx2.1*-lineage neurons, including about 30% of cortical interneurons, in cKO mice^15^. Combined with the known role of cortico-septal connections in the regulation of certain behaviors^19,63^, this raises the possibility that part of the behavioral phenotype we observed in cKO animals could be due to circuit disruptions in other parts of the brain beyond the LS. For this reason, we used a forced restraint assay to obtain direct evidence of the activation of LS *Nkx2.1*-lineage neurons upon exposure to a stressful stimulus. As expected, we found a robust and consistent increase in the density of c-Fos+ neurons in the lateral septum of restrained mice^56,65–67^, which was lower in cKO mice, particularly in the LSd. The proportion of activated *Nkx2.1*-lineage neurons was approximately doubled in restrained animals compared to controls, further confirming the involvement of this population in stress responses (**Figures 6, S6**). However, future experiments directly measuring and/or specifically manipulating the activity of *Nkx2.1*-lineage neurons in the LS will be necessary to refine our knowledge about their involvement in maintaining the balance between threat avoidance and novelty seeking.

Taken together, our results fit with the current view of the lateral septum as a key component in the calibration of behavioral responses based on the motivational state and the external context of the animal^2,3^. Our data provides new insight into the molecular identity and behavioral significance of the *Nkx2.1*-lineage/*Crhr2*-expressing LS neuronal population, establishing a novel model where it is almost entirely ablated. Our study highlights how understanding the developmental origin of different neuronal subtypes in the lateral septum can be an invaluable step towards understanding their mature identity and function.

## METHODS

### EXPERIMENTAL MODEL AND SUBJECT DETAILS

All animal procedures conducted in this study followed experimental protocols approved by the Institutional Animal Care and Use Committees of Harvard Medical School (HMS) and the University of California, San Francisco (UCSF). Mouse strains mentioned in the main text are listed in the **Key Resources** table. Mouse housing and husbandry were performed in accordance with the standards of the Harvard Medical School Center of Comparative Medicine (HCCM) at HMS, and the Laboratory Animal Resource Center (LARC) at UCSF. Mice were group housed in a 12 h light/dark cycle, with access to food and water *ad libitum*. Samples were obtained at the ages indicated in the Figure Legends and throughout the text. Results reported include animals of both sexes, balanced equally whenever possible, except where otherwise indicated.

## METHOD DETAILS

### Tissue processing for histology

Mice were transcardially perfused with PBS followed by 4 % paraformaldehyde (PFA) in 120 mM phosphate buffer; their brains were dissected out and post-fixed in 4 % PFA overnight at 4°C. Brains were sectioned into 50-100 µm sections on a vibratome and stored at -20°C in freezing buffer (30 % ethylene glycol, 28 % sucrose in 0.1 M sodium phosphate buffer), or at 4°C in PBS with 0.05 % sodium azide.

### Immunofluorescence

Floating vibratome sections (50 µm thick) were permeabilized with 0.5 % Triton X-100 in PBS for 1-2 h, then blocked with blocking buffer (10 % goat serum, 0.1 % Triton X-100 in PBS) for 1-2 h at room temperature. They were then incubated for 24-72 h, at 4°C, with primary antibodies diluted (see **Key Resources** table for dilutions) in blocking buffer. The samples were washed three times (10-30 min/wash) with PBS, counterstained with DAPI (4’,6-diamidino-phenylindole) for 45 min (both steps at room temperature), and incubated with secondary antibodies diluted in blocking buffer for 2 h at room temperature or overnight at 4°C. They were then washed (three 10-30 min washes) with PBS, and mounted on microscope slides with ProLong Gold Antifade Mountant.

### Microscopy and image analysis

Images were acquired using either a Leica Stellaris laser point scanning confocal microscope or a Leica DM5500B widefield microscope. 10x, 20x and 40x objectives were used, and the parameters of image acquisition (laser intensity, speed, resolution, averaging, zoom, z-stack, etc.) were adjusted for each set of samples. Images were further analyzed using ImageJ, both in its native and Fiji distributions, as described below. Brightness and contrast were adjusted as necessary for visualization; the source images were kept unmodified.

#### Cell quantification

All cell type quantifications were done with the CellCounter tool in ImageJ/Fiji. All cells positive for the corresponding marker(s) within the dorsal, intermediate and ventral nuclei of the lateral septum (LSd, LSi, and LSv, respectively) were counted; values reported are the average across both hemispheres for each sample. For the quantifications in **Figures 1E and S1G**, the medial septal nucleus (MS) was defined as shown in **Figure 1B** (i.e., with its ventral extent reaching as far as that of the lateral ventricles), and all cells within it were counted. The rostro-caudal locations labeled as ‘I’, ‘II’, and ‘III’ throughout the manuscript correspond approximately to Bregma +0.75, +0.5 and +0.25, respectively.

#### Septum size measurement

For **Figures 1F and S1D**, 50- to 70-µm vibratome-sliced coronal sections corresponding to rostro-caudal sections I-III were measured. All sections had been subjected to immunofluorescence staining for other figures. The total area of the section and the area of the septum (defined as the part dorsal to a straight line drawn between the bottom corners of both lateral ventricles) were calculated in Fiji after manually tracing them in the DAPI channel. The average proportion of septum/total section area of WT brains was used to normalize the measurements.

#### Fluorescence intensity quantification

For **Figures 3C and S3C**, the mean fluorescence intensity was calculated across the entirety of each area analyzed, using the ‘Measure’ tool in ImageJ, and then normalized by the background fluorescence level, which was calculated as the mean intensity of a 50 x 50 µm square located within the same area that presented no obvious urocortin-3 (**Figure 3A**) or enkephalin (**Figure S3A**) staining.

#### Perineuronal basket quantification

For **Figures 3D-F and S3D**, baskets were only quantified as such when the signal consistently surrounded at least 50 % of the perimeter of a neuron with visible DAPI counterstaining within the presumed soma. Examples are highlighted in **Figure 3B**.

### Fluorescent *in situ* hybridization

50-µm sections (obtained as described above) were placed on slides and submitted to the RNAscope protocol (Advanced Cell Diagnostics), following the manufacturers’ instructions with minor modifications. The RNAscope probe (listed in the **Key Resources** table) was purchased from ACD. In **Figures 2I-K** and **S2G-I**, cells were quantified as positive for *Crhr2* if there were five or more RNAscope puncta in their nucleus and/or soma.

### Single-nucleus RNA-sequencing and library preparation

Mice were sacrificed at postnatal day 35. Their septa were manually dissected in hibernation buffer (0.25 M sucrose, 25 mM KCl, 5 mM MgCl_2_, 20 mM Tricine KOH) on ice, and transferred to a pre-chilled 2 ml Dounce homogenizer containing 500 µl of dissociation buffer (hibernation buffer supplemented with 5 % IGEPAL, half a Roche protease inhibitor tablet, 0.2 U/µl Promega RNasin, 0.1 % spermidine, 0.1 % spermine, and 0.1 % dithiothreitol [DTT]). The tissue was triturated with a loose pestle followed by a tight one, performing ∼15 strokes with each. After resting on ice for 10 min, the homogenate was transferred to a low-bind Eppendorf tube and centrifuged for 5 min at 500xg, at 4°C. The supernatant was discarded, and the pellet was gently resuspended in 500 µl of PBS containing 1 % bovine serum albumin, 0.2 U/µl Promega RNAsin, 0.1 % spermidine, 0.1 % spermine, and 0.1 % DTT. The centrifugation and pellet resuspension steps were repeated once before straining the resulting suspension through a 40 µm Flowmi filter and staining with 0.8 µl of Ruby Dye. Ruby Dye-positive nuclei were sorted into GFP-positive and GFP-negative populations using a Sony SH800S Fluorescence Activated Cell Sorter, collecting a minimum of 50,000 events of either population. Nuclei were then loaded onto a 10X Chromium Single Cell 3’ chip (v3 chemistry), at optimal concentrations as recommended by the manufacturer, and submitted to the system’s standard protocol. The resulting libraries were sequenced using a NovaSeq 6000 S4 flow cell, at a read depth of ∼50,000 reads/nucleus. A total of 12 mice (3 males and 3 females each for WT and cKO conditions), across six replicates, were used. The WT samples reported here are the same ones used in Reid *et al*. 2024^24^.

### Single nucleus RNA-sequencing analysis

Sequenced reads were aligned to the mouse genome (assembly mm10), including intronic reads, using the Cell Ranger 7.1.0 software package from 10X Genomics. All subsequent analyses were performed in R Studio, using R version 4.3.3, using the Seurat package version 5.2.0 (https://satijalab.org/seurat/). Genes expressed in fewer than 3 nuclei were excluded from further analyses. Nuclei expressing fewer than 200 or more than 7,000 genes, and/or fewer than 1,000 or more than 30,000 RNA counts, and/or more than 0.5 % mitochondrial RNA counts were filtered out, resulting in a total of 24,323 nuclei (**Figure S2A**). Raw counts were log-normalized to 10,000 reads per cell, and the top 2,000 highly variable genes were identified. After performing principal component analysis (PCA), a neighborhood graph was constructed using the first 10 principal components, and subsequently reduced to two dimensions using Uniform Manifold Approximation and Projection (UMAP) for visualization and exploration of the data. Nuclei were clustered at a resolution of 0.5 for **Figure S2A**, and the expression of known marker genes (**Figure S2B**) was used to select clusters 0, 1, and 4 as the subset of the initial dataset containing septal neurons. This subset was analyzed in a similar way (neighborhood graph with 30 PCs, UMAP clustering at 0.8 resolution). Based on marker gene expression, cluster 13 of the resulting dataset was excluded as a non-septal population, and the analysis was reiterated for the remaining 21 clusters, obtaining a septal neuron dataset composed of 8,783 nuclei (**Figure S2C**). This was further subdivided into medial and lateral septum neurons based on the expression of select marker genes (**Figure S2D**). The final lateral septal neuron dataset (4,984 nuclei; **Figure 2B**) was composed of clusters 0, 1, 6, 7, 8, 9, 11, 13, 15, 16, 17 and 18 from the total septal neuron dataset; it was subsequently analyzed using the same parameters as above to define the different LS neuronal populations.

### Electrophysiology recordings

Mice (P18-P23) were deeply anesthetized with isoflurane and decapitated. Their brains were removed and placed in ice-cold modified artificial cerebrospinal fluid (ACSF) composed of (in mM): 87 NaCl, 26 NaHCO_3_, 2.5 KCl, 1.25 NaH_2_PO_4_, 0.5 CaCl_2_, 4 MgCl_2_, 10 glucose, 75 sucrose, saturated with 95% O_2_/5% CO_2_, and with pH = 7.4. A vibratome was used to obtain coronal slices, which were incubated at 34°C for 30 min and stored at room temperature until used. They were then transferred to the recording chamber of a Zeiss Axioskop microscope, where they were continually perfused with ACSF composed of (in mM): 124 NaCl, 24 NaHCO_3_, 2.5 KCl, 1.2 NaH_2_PO_4_, 2 CaCl_2_, 2 MgSO_4_, 5 HEPES, 12.5 glucose (300-310 mOsm, pH = 7.4), and maintained at room temperature during recordings. Recording pipettes were made of pulled borosilicate glass capillaries with a resistance of 3-5 MΩ when filled with internal solution (in mM: 130 K D-Gluconate, 5 KCl, 10 HEPES, 0.2M EGTA.KOH, 8 Phosphocreatine Tris_2_, 4 MgATP, 0.2 Na_2_GTP, 0.4% Biocytin, pH 7.2-7.4 and 295-305 mOsm/l). Cells were visualized using a 40x IR water immersion objective; whole-cell recordings were obtained from randomly selected LS neurons positive or negative for tdTomato, from either WT or cKO animals, as indicated in the figure panels and legends. Recordings were performed with a Multiclamp 700B amplifier and digitized using a Digidata 1440A and the Clampex 10 program suite. N = 6 (4 males, 2 females) for WT samples; N = 5 (all males) for cKO. The numbers of cells analyzed for each measurement (n) are indicated in the corresponding figure legends.

### Behavior

#### Anxiety-related tests

A series of tests were carried out to assess general anxiety-like behavior in WT *vs*. cKO mice. They were performed as described below, in the order laid out in **Figure S5A**: after an initial 2-week period of acclimation to the facilities of the Mouse Behavior Core at Harvard Medical School, mice were subjected to the light-dark box, open field and novel object tests, with one day in between tests; after 5 days to ten days, they were subjected to the elevated plus-maze and social interaction tests, with one day between both tests and another between the habituation and test portions of the second one; finally, after another period of one week to ten days, they were subjected to the Y-maze test. Experiments were conducted in three separate batches of 2-6 months old animals including WT, cKO, and heterozygote controls (Nkx2.1-Cre;Prdm16^+/f^;Ai14+), from multiple litters. All mice were socially housed, with a minimum of 2 and a maximum of 5 littermates per cage. Male and female mice were tested separately, and the experimenters were blinded to the animals’ genotypes until after the results were compiled. The experimental arenas were thoroughly cleaned (successively wiped with Windex and 70% ethanol) between individual tests.

##### Light-dark box test

The arena (open field chamber [ENV-510] with a dark box insert [ENV-511]; Med Associates) consisted of two equally-sized compartments connected by a small restricted opening. One of the compartments was brightly illuminated (700-800 lux), while the other remained in the dark. Mice were placed on the bright compartment and allowed to freely move within and between both chambers for 10 minutes. The number of entries and time spent in each compartment were recorded automatically using Activity Software (Med Associates). The measure compared in **Figure 5C** is % of the test time spent in the bright chamber.

##### Open field test

The arena was a uniformly illuminated box with a square (40×40 cm) base and an open top. Mice were placed in a corner of the box and allowed to freely move within it for 10 minutes. Their movements were recorded with a camera placed above the center of the arena, and analyzed using the Topscan software (CleverSys). The measure compared in **Figure 5A** is % of the test time spent in the center of the arena.

##### Novel object test

This test was performed and analyzed in the same arena and manner as the open field test, the only difference being the placement of a novel object (a small assembly of Lego pieces, approximately 4×4×7 cm in size) attached to the center of the base. The measure compared in **Figure 5B** is % of the test time spent in the middle of the arena.

##### Elevated plus-maze test

The arena consisted of two open and two (laterally) closed arms, all measuring 30 x 5 cm and extending from a central platform, located 85 cm above ground level. Mice were placed at the end of one of the open arms, facing the center, and allowed to freely explore the arena for 5 minutes in a dimly lit environment. Their movements were recorded with a camera placed above the center of the maze, and analyzed using the Topscan software (CleverSys). The measure compared in **Figure S5B** is % of the test time spent on the open arms.

##### Social interaction test

This test was performed on a three-chambered rectangular (62 x 40 cm) arena. The central chamber was separated from the side ones by restricted openings that could be blocked by removable doors. Each side chamber had a small perforated container located on a corner (in opposite locations). The test was performed in two phases, on two consecutive days. In the initial (habituation) phase, the test mouse was placed in the middle chamber, the doors to the side chambers were removed simultaneously and the subject was allowed to freely explore the arena for 10 minutes. In the second (test) phase, a “stranger” mouse (C57BL/6J strain, of the same sex as the test subject and with no prior contact with it) was placed under one of the perforated containers in the side chambers, and an object (the same Lego assembly that was used for the novel object test) under the other, in a random and counterbalanced way. Test mice were placed in the middle chamber and allowed to freely explore the arena for 10 minutes after removing the doors to the side chambers. Their movements were recorded and analyzed with the Topscan software (CleverSys). The measure compared in **Figure S5C** is the proportion of time spent in close proximity (within 2.5 cm) to the container where the unfamiliar mouse was placed, over the total amount of time spent exploring (i.e. close to) either container.

##### Y-maze test

This test was performed in a transparent Y-shaped arena enclosed on the sides, where two arms (“left” and “right”) can be independently blockaded, with the third one serving as the starting arm. The arena was placed close to a large green holding rack, which served as a visual spatial cue for mice to differentiate the sides of the testing room. The test was run in two phases: during the habituation phase, either the left or the right arm was blockaded, and mice were placed at the end of the starting arm. They were allowed to freely explore the arena for 3 minutes, while their movements were recorded. They were then retrieved from the arena and placed in a designated empty cage, while the arena was cleaned and the blockade removed. During the test phase, the mice were put back into the maze in the same manner, and allowed to explore the arena for 3 minutes. Their movements were recorded and analyzed using the TopScan software (CleverSys). The measure compared in **Figure S5D** is the proportion of time spent on the novel arm (i.e., the one that had been blockaded during the habituation phase) over the total amount of time spent on either “left” or “right” arms.

#### Acute restraint stress

Mice (6-7 months old) were placed inside 40 ml Falcon tubes with ventilation holes bored in at the tip (i.e. where the mouse’s head was located). The tube caps were screwed in and the tubes were placed in a designated clean cage for 30 minutes. Mice were then removed from the tubes into their home cages, and sacrificed by transcardial perfusion 1 hour later. Male and female mice were subjected to restraint on different days, and the tubes were thoroughly cleaned with 70% ethanol and dry-wiped between subjects. Untreated age-matched controls were submitted to perfusion as described above, without any further experimental manipulations.

#### Predator Odor Exposure

A circular open field (US Plastics) with a small l1” diameter odor inlet on one side of the arena was used to expose animals for 15-minutes to a “blank” odor (dipropylene glycol), then 15-minutes of a 25% dilution of 2,3,5-trimethyl-3-thiazoline (TMT; diluted in dipropylene glycol). Filtered air (0.2-0.25 liters per minute) was blown over odorant-soaked filter paper (VWR) placed at the bottom of a Vacutainer syringe vial (Covidien). Independent PTFE tubing lines (Zeus) were used for blank and TMT sessions to avoid cross-contamination. All recordings were performed in darkness. The arena was cleaned with successive 70% ethanol – 10% bleach – 70% ethanol wiping steps and allowed to dry completely between subjects. Male and female mice were tested on different days.

##### Occupancy Analysis

Heatmaps (**Figure 5E**) were generated using the (x,y) centroid position of animals calculated during the extraction procedure. Specifically, a 2D histogram was generated using all data from both male and female datasets using the numpy (python) histogram2d() function with a bin size of 15. The specific x and y bin edges from this “grand” heatmap were used to generate all subsequent groupwise heatmaps. To calculate the average heatmap for a given condition, a 2D histogram was generated for each session, normalized so that the histogram summed to 1, then appended to a (n_sessions x 15 bins x 15 bins) matrix. Finally, the average was taken across the first dimension to create a final (15×15) dimensional matrix. For heatmaps where differential occupancy is shown, average (15×15) heatmaps were simply subtracted from each other. For visualization purposes, all heatmaps were smoothed with a gaussian kernel using the matplotlib imshow() function. Odor inlet occupancy was quantified by computing the distance (vector norm) between the mouse centroid and a fixed odor port position, then counting the proportion of time the mouse spent closer to the odor port than a given threshold (40 mm in **Figure 5D,F**). Outliers (a total of four, one per sex and genotype) were identified by the Grubbs’ test, performed online with the GraphPad Outlier calculator at https://www.graphpad.com/quickcalcs/grubbs1/, and excluded from statistical analyses.

#### Motion Sequencing (MoSeq)

##### Data Acquisition

Motion Sequencing (**Figures S5E-L**) is an unsupervised machine learning platform that segments continuous mouse behavior into its constituent parts, or behavioral “syllables”^42,68–70^. Data acquisition was performed as previously described^71^. Briefly, we used “depth” MoSeq, which operates on 3D data acquired by a depth sensor. Specifically, kinect2-nidaq software (https://github.com/dattalab/kinect2-nidaq) was used to acquire 3D images from a Kinect 2 depth camera at 30 Hz.

##### MoSeq: Pre-processing, Data Extraction, Modeling

Data was extracted, pre-processed, and modeled using the moseq2-app pipeline (https://github.com/dattalab/moseq2-app). Male and female mice used in this study had drastically different distributions of sizes; since MoSeq is sensitive to deviations of this magnitude, we separated male (1,669,346 frames) and female (1,346,820 frames) datasets for subsequent MoSeq analysis. In addition, the top/bottom 2.5% sized animals (mean mouse area) were omitted from each dataset, as detailed previously^72^. Default parameters were used for extraction and PCA. To determine an ideal value of the hyperparameter kappa, a scan over a log range of 10 kappa values was performed on each dataset and a 100-iteration AR-HMM model was trained for each value of kappa. Following the kappa scan, the best kappa value (1,668,101 for both datasets) was selected using the Jensen-Shannon objective, which measures the similarity between the AR-HMM learned syllable duration distribution and the model-free changepoint duration distribution. A final 1000-iteration model was trained with a kappa of 1,668,101. The final model for each dataset had an overlapping syllable duration distribution and changepoint distribution similar to previously published datasets. All pre-processing, data extraction, and modeling were performed locally on a Windows 10 workstation.

##### MoSeq: Analysis

The moseq2-app pipeline was also used for syllable analysis. Syllables that accounted for 99% of variance explained were used for downstream analysis (53 for males, 53 for females). Using the interactive syllable labeling tool, syllables were given human labels by examining crowd movies. Differential syllable usage analysis was performed on all pairwise group comparisons for each dataset. Significantly different syllable usages were determined using a Kruskal-Wallis test, post-hoc Dunn’s two-sided test with permutation, and multiple comparisons correction using the Benjamini-Hochberg procedure with a false discovery rate of 0.05^43^. All syllable usage error bars report the 95% confidence interval from 10,000-iteration bootstrapped syllable usages.

### Statistical analysis

All statistical analyses performed are indicated in the corresponding Figure Legends. Except for MoSeq (see above), they were performed using GraphPad Prism 9. All p-values were rounded to ten thousandth, and are presented above each statistical comparison in the corresponding figures; p-values below 0.05 (which we considered the cutoff for statistical significance) are highlighted in bold, while p-values deemed not statistically significant under this criterion are displayed in regular type within parentheses.

## KEY RESOURCES TABLE

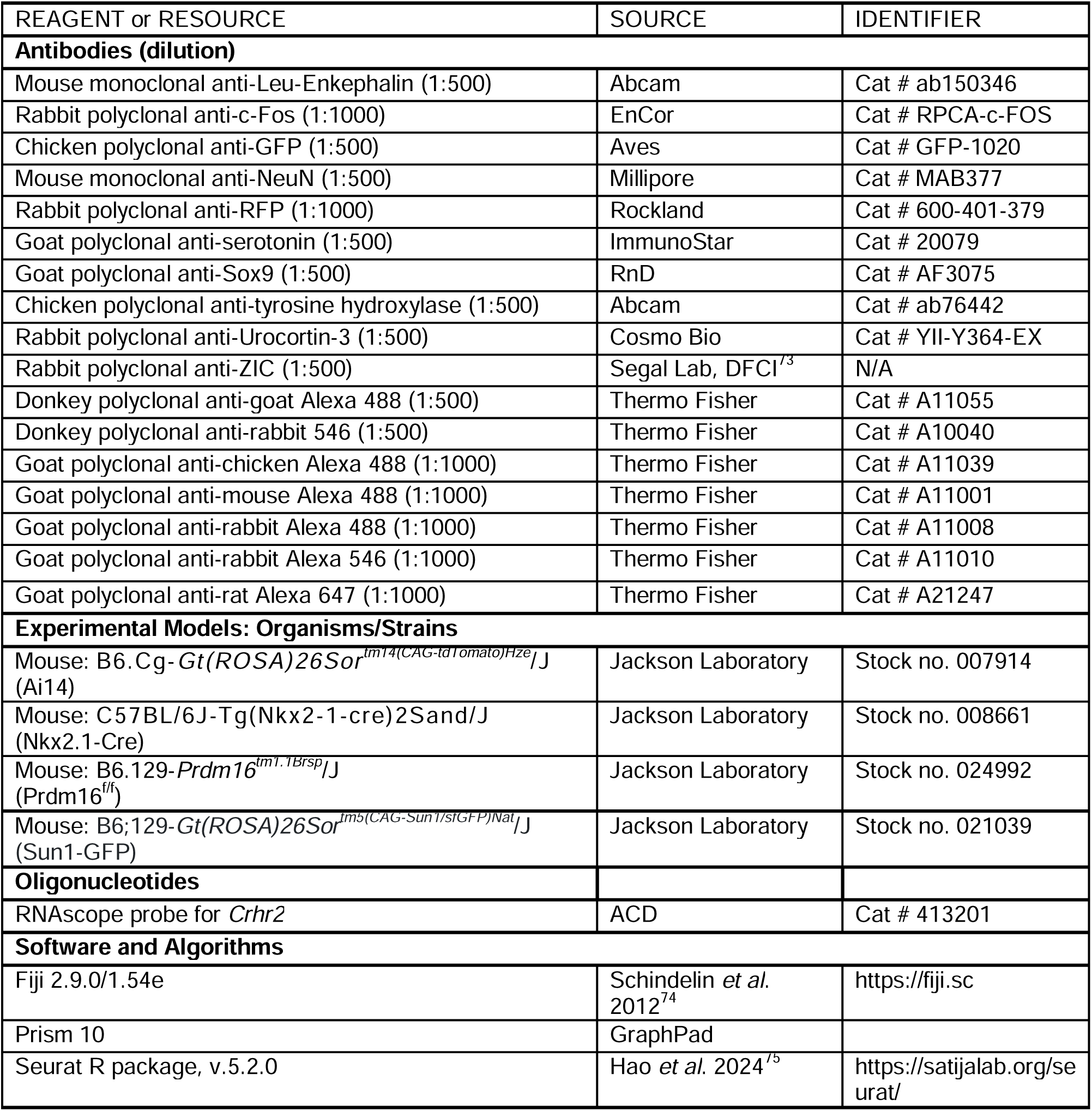

## ACKNOWLEDGEMENTS

The authors would like to thank all members of the Harwell lab for feedback and support; Manni Adam, Rhiana Simon, Félix Leroy and Antoine Besnard for comments on the manuscript; Barbara Caldarone at the Mouse Behavior Core of Harvard Medical School for her help and advice towards the design, performance and analysis of anxiety-related behavioral tests; Bob Datta for providing advice, equipment and resources for the predator odor assay; Mazen Kheirbek for providing equipment and laboratory space for the acute restraint tests; Maria Pazyra and Rosalind Segal for their gift of anti-ZIC antibody; and members of the Álvarez-Buylla, Canzio, Goodrich, Kriegstein, Lehtinen, Panagiotakos, Paredes, and Segal labs for discussions and feedback. M.T.G. was partially supported by the Ellen R. and Melvin J. Gordon Center the Cure and Treatment of Paralysis. This research was funded by grants R01NS102228 and R01MH119156 from the National Institutes of Health.

## AUTHOR CONTRIBUTIONS

Conceptualization: M.T.G., C.C.H.; Software: R.E.P.; Formal analysis: M.T.G., R.E.P., L.A.I., Y.R.; Investigation: M.T.G., D.N.T., R.E.P., L.A.I., C.M.R., S.M.H., F.D-H, S.K.S., Y.X., S.V.; Resources: C.C.H.; Data curation: M.T.G., R.E.P.; Writing – original draft preparation: M.T.G.; Writing – review and editing: M.T.G., D.N.T., S.K.S., C.M.R., Y.X., C.C.H.; Visualization: M.T.G., R.E.P.; Supervision: M.T.G., C.C.H.; Funding acquisition: C.C.H.

## SUPPLEMENTARY FIGURE LEGENDS

**Figure S1.**
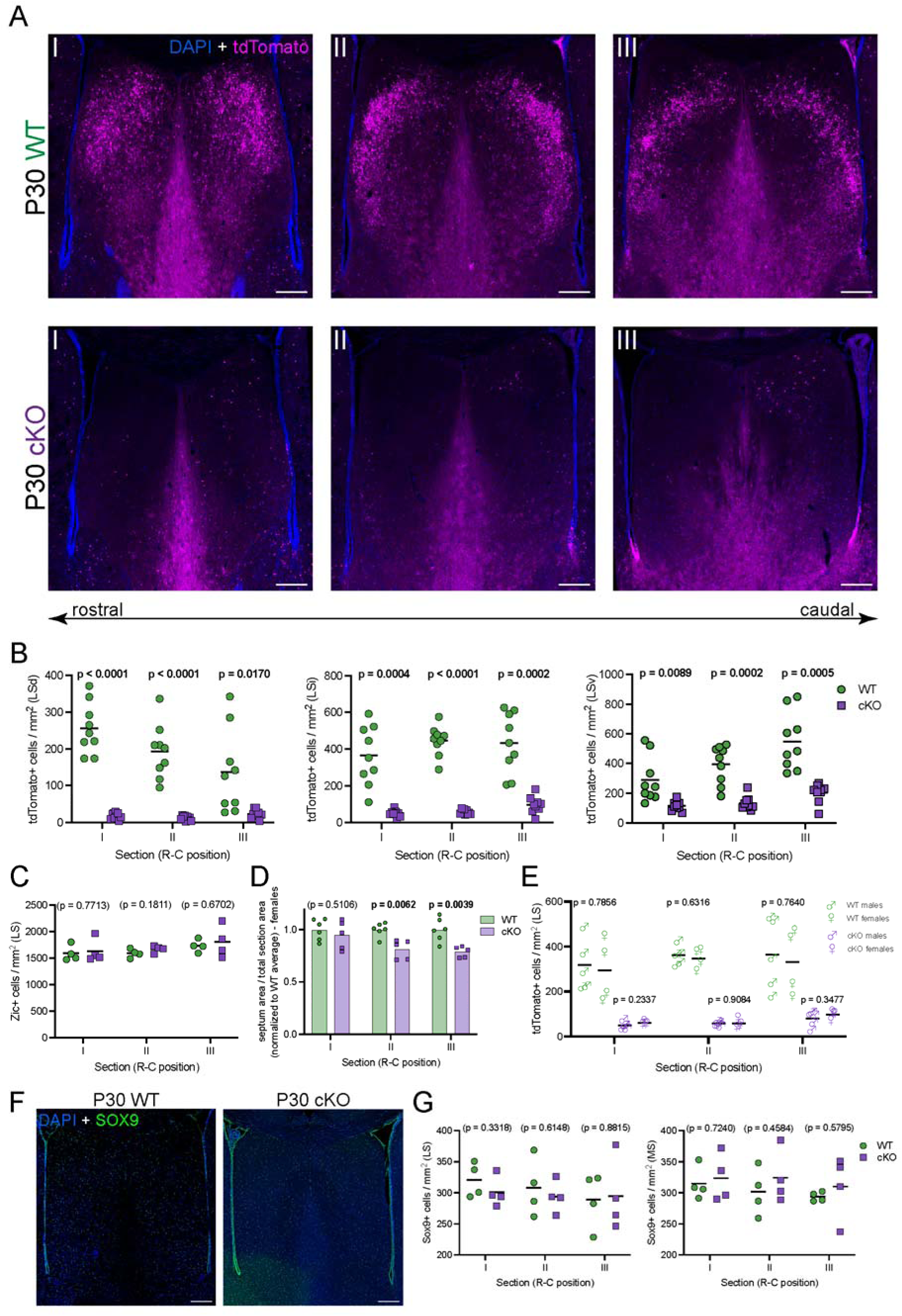
related to Figure 1. **A)** Coronal sections along the rostral (left) to caudal (right) axis through the forebrain of P30 WT (top) and cKO (bottom) mice, submitted to immunofluorescence staining for tdTomato (magenta) and counterstained with DAPI (blue). Scale bars, 250 µm. **B)** Quantification of the density of cells positive for tdTomato per mm^2^ in the different subnuclei within the lateral septum of WT (green circles, n = 9) and cKO (purple squares, n = 9) mice. LSd, dorsal lateral septum; LSi, intermediate lateral septum; LSv, ventral lateral septum. **C)** Quantification of the density of cells positive for Zic per mm^2^ in the lateral septum of WT (green circles, n = 4) and cKO (purple squares, n = 4) mice. **D)** Quantification of the area of the septum relative to the total area of its corresponding coronal brain section in WT (green circles, n = 6) and cKO (purple squares, n = 5) female mice. Measurements are normalized to the corresponding WT average. **E)** Quantification of the density of cells positive for tdTomato per mm^2^ in the lateral septum of WT (green symbols, n = 5 males, 4 females) and cKO (purple symbols, n = 5 males, 4 females) mice, as in **Figure 1D**, separated by sex as indicated. **F)** Example images of the septa of P30 WT (left) and cKO (right) mice, submitted to immunofluorescence staining for Sox9 (green) and counterstained with DAPI (blue). Scale bars, 250 µm. **G)** Quantification of the density of cells positive for Sox9 per mm^2^ in the lateral (left) and medial (right) septum of WT (green circles, n = 4) and cKO (purple squares, n = 4) mice. Unpaired t-tests with Welch’s correction were performed; the p-values are shown above the corresponding compared sets of data: bold typeface indicates statistically significant (p<0.05) differences.

**Figure S2.**
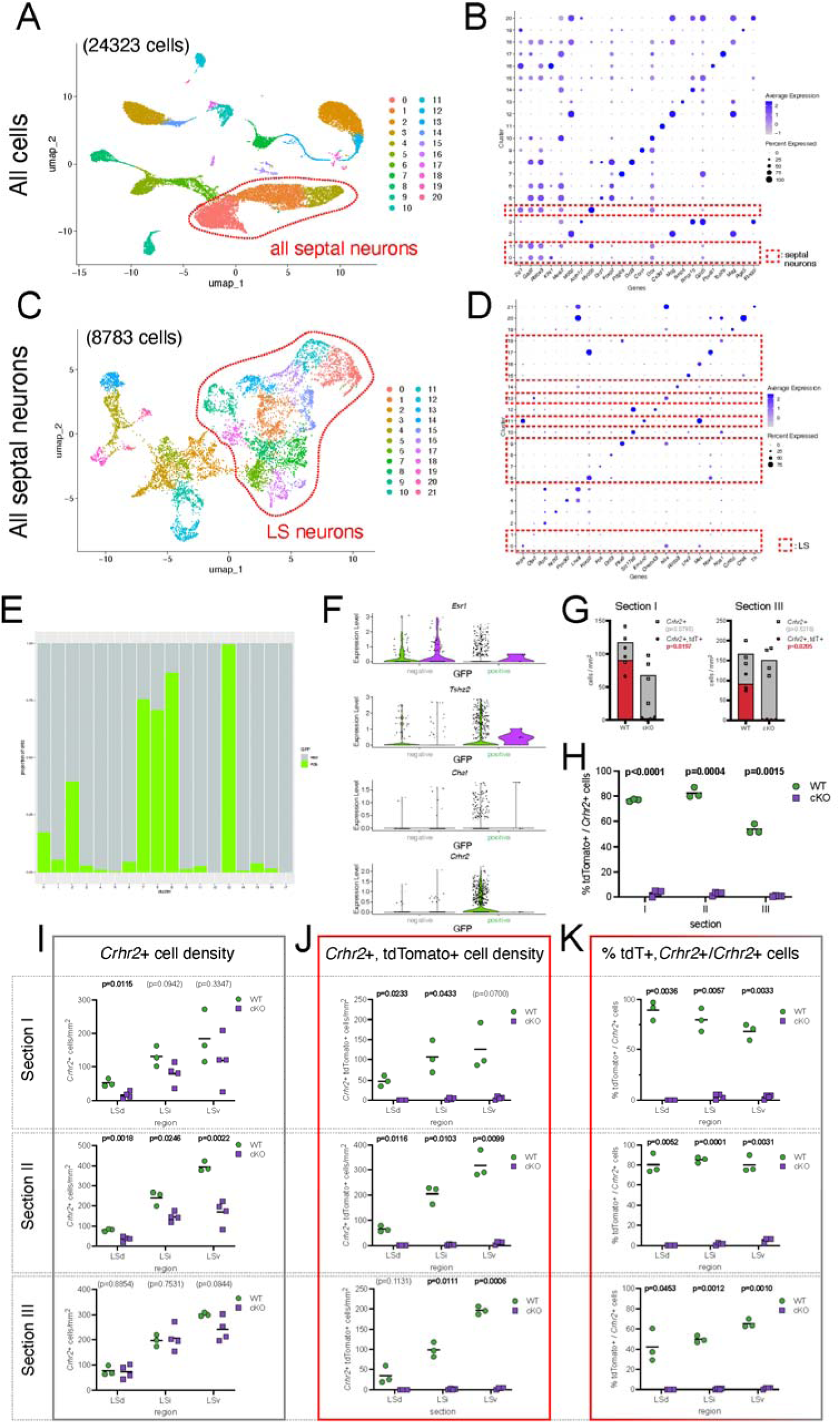
related to Figure 2. **A)** Uniform Manifold Approximation and Projection (UMAP) plot representing a 2D dimensional reduction of the transcriptional similarity among all cells identified within the dataset (24323 cells from a total of 6 WT and 6 cKO animals [3 males and 3 females each]). **B)** Dot plot showing the expression of select marker genes in each of the clusters in **A**, allowing their identification as distinct cell types. Clusters containing septal neurons are highlighted by dashed red lines. **C)** UMAP plot of all septal neurons as identified in **A** and **B** (8783 cells). **D)** Dot plot showing the expression of select marker genes in each of the clusters in **C**, allowing their identification as distinct cell types. Clusters containing lateral septum neurons are highlighted by dashed red lines. **E)** Bar graph showing the proportion of cells belonging to the Nkx2.1-lineage (green) *vs*. all other cells (gray) within each of the 17 clusters outlined in **Figure 2B**. **F)** Violin plots showing the expression levels of select marker genes within clusters 7, 8, 9, and 13 of **Figure 2B** (i.e. all clusters where ≥75% of cells belong to WT samples), split by genotype (WT, green; cKO, purple) and lineage (Nkx2.1-lineage: GFP-positive; all other lineages: GFP-negative). **G)** Bar graphs showing the density (per mm^2^) of all cells positive for *Crhr2* (gray squares) and the subset of those cells that are also tdTomato+ (red circles), in sections I and III of WT *vs*. cKO animals. **H)** Quantification of the percentage of *Crhr2*+ cells that are tdTomato+ throughout the entire LS of WT (green circles) *vs*. cKO (purple squares) animals. **I-K)** Detailed quantifications of total *Crhr2*+ cell density (**I**); *Crhr2*+, tdTomato+ cell density (**J**); and % of tdTomato+ cells within the *Crhr2*+ population (**K**), split by sections across the rostro-caudal axis (sections I, II and III) and by LS subdivisions (LSd, LSi, LSv), as shown in **Figure 1**. **G-K:** (N = 3 [2 males, 1 female] for WT; N = 4 [2 males, 2 females] for cKO)

**Figure S3.**
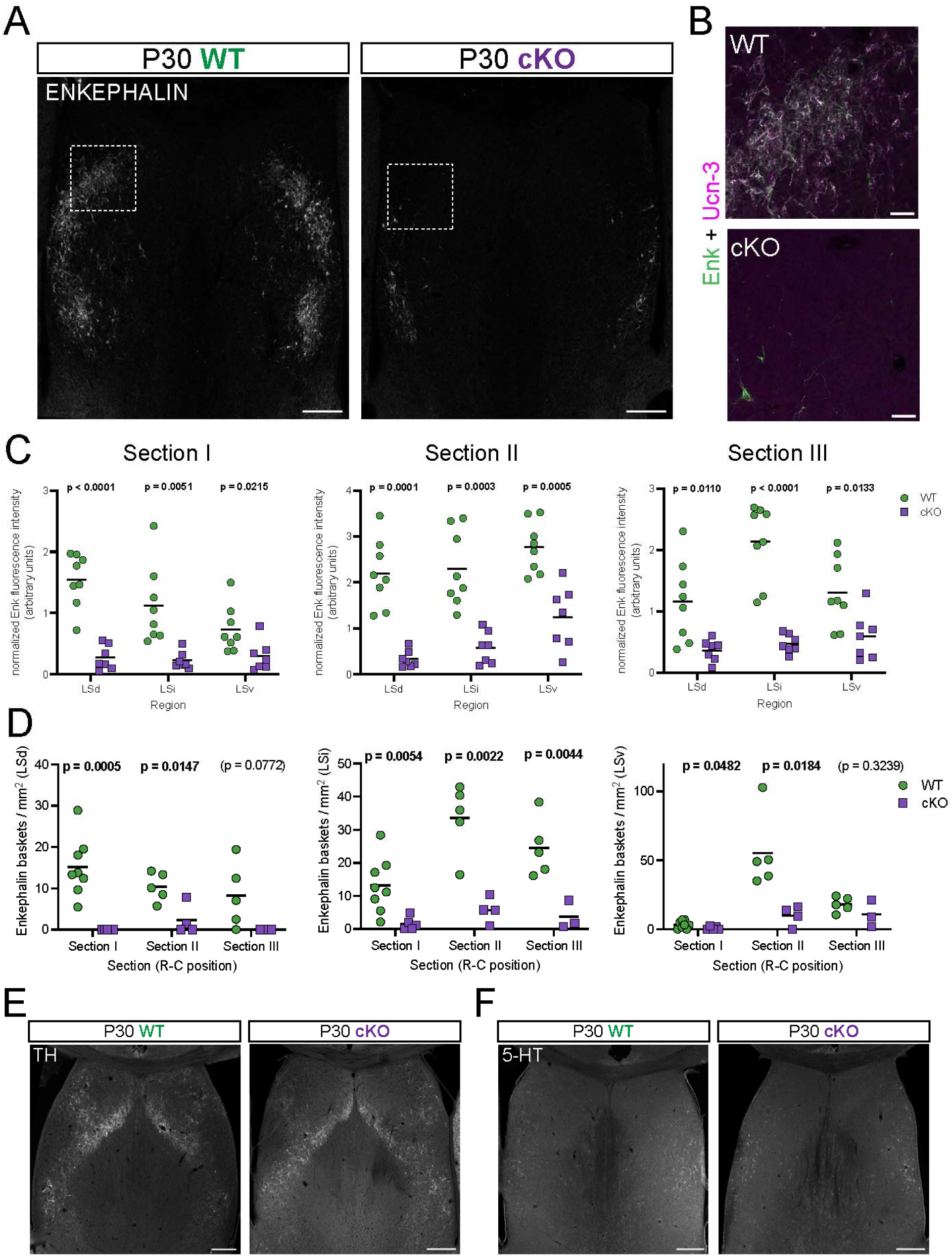
related to Figure 3. **A)** Overview coronal images of the septum of P30 WT (left) and cKO (right) mice submitted to immunofluorescence staining for enkephalin (gray). Scale bars, 250 µm. **B)** Closeup view of the white dashed line boxes in **A**, showing the combined signals for enkephalin (green) and urocortin-3 (magenta), in the LS of WT (top) and cKO (bottom) mice. Scale bars, 50 µm. **C)** Quantification of fluorescence intensity in the different LS subnuclei of P30 WT (green circles) and cKO (purple squares) brains where enkephalin was detected by immunofluorescence staining, at three levels along the rostro-caudal axis (Sections I through III). **D)** Quantification of the density of enkephalin+ baskets per mm^2^ in the different LS subnuclei, at three levels along the rostro-caudal axis (Sections I through III), in WT (green circles) and cKO (purple squares) mice. **E)** Overview coronal images of the septum of P30 WT (left) and cKO (right) mice submitted to immunofluorescence staining for tyrosine hydroxilase (TH, gray). Scale bars, 250 µm. **F)** Overview coronal images of the septum of P30 WT (left) and cKO (right) mice submitted to immunofluorescence staining for serotonin (5-HT, gray). Scale bars, 250 µm. **C-D:** N = 8, N = 5 and N = 5 for WT; N = 5, N = 4 and N = 3 for cKO in Sections I, II, and III, respectively. Unpaired t-tests with Welch’s correction were performed; the p-values are shown above the corresponding compared sets of data: bold typeface indicates statistically significant (p<0.05) differences.

**Figure S4.**
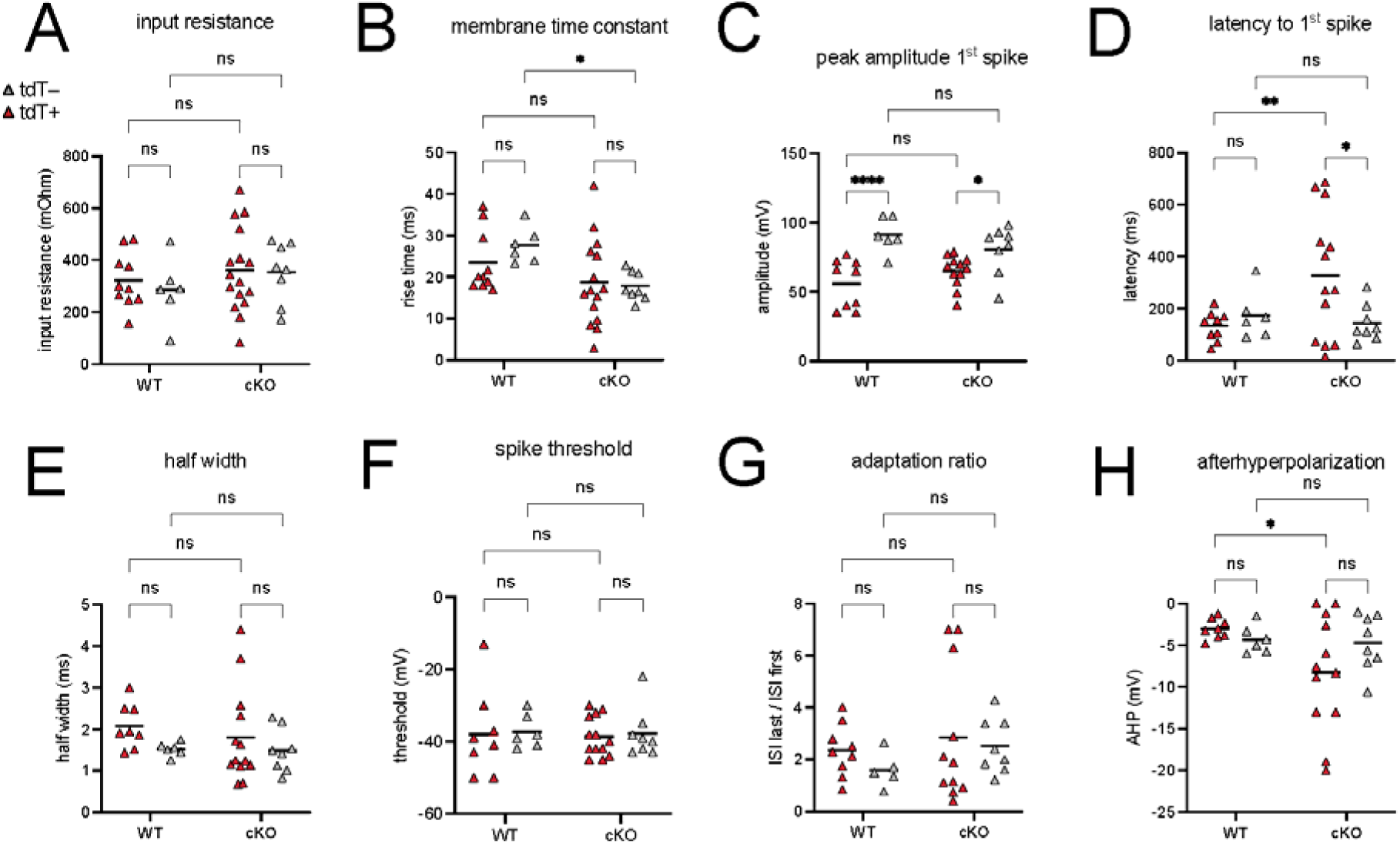
**A-H)** Comparison of intrinsic electrophysiological properties of tdTomato+ and tdTomato– neurons in the LS of WT and cKO samples at 3 weeks of age: **A**, input resistance; **B**, membrane time constant; **C**, peak amplitude of the first spike; **D**, latency to the first spike; **E**, half-width; **F**, spike threshold; **G**, adaptation ratio; **H**, afterhyperpolarization. After removal of outliers via Grubbs’ test, 2-way ANOVA tests with multiple comparisons were performed; the p-values are shown above the corresponding compared sets of data: bold typeface indicates statistically significant (p<0.05) differences. For WT tdTomato+, n = 10 (A, B), n = 9 (C, D, G), n = 8 (E, F, H); for WT tdTomato–, n = 6 (A, B, C, D, E, F, H), n = 5 (G); for cKO tdTomato+, n = 16 (A,) E, G, J, K, Q, R), n = 15 (B), n = 13 (D, E, F), n = 12 (C, H), n = 11 (G); for cKO tdTomato–, n = 8 (A-H).

**Figure S5.**
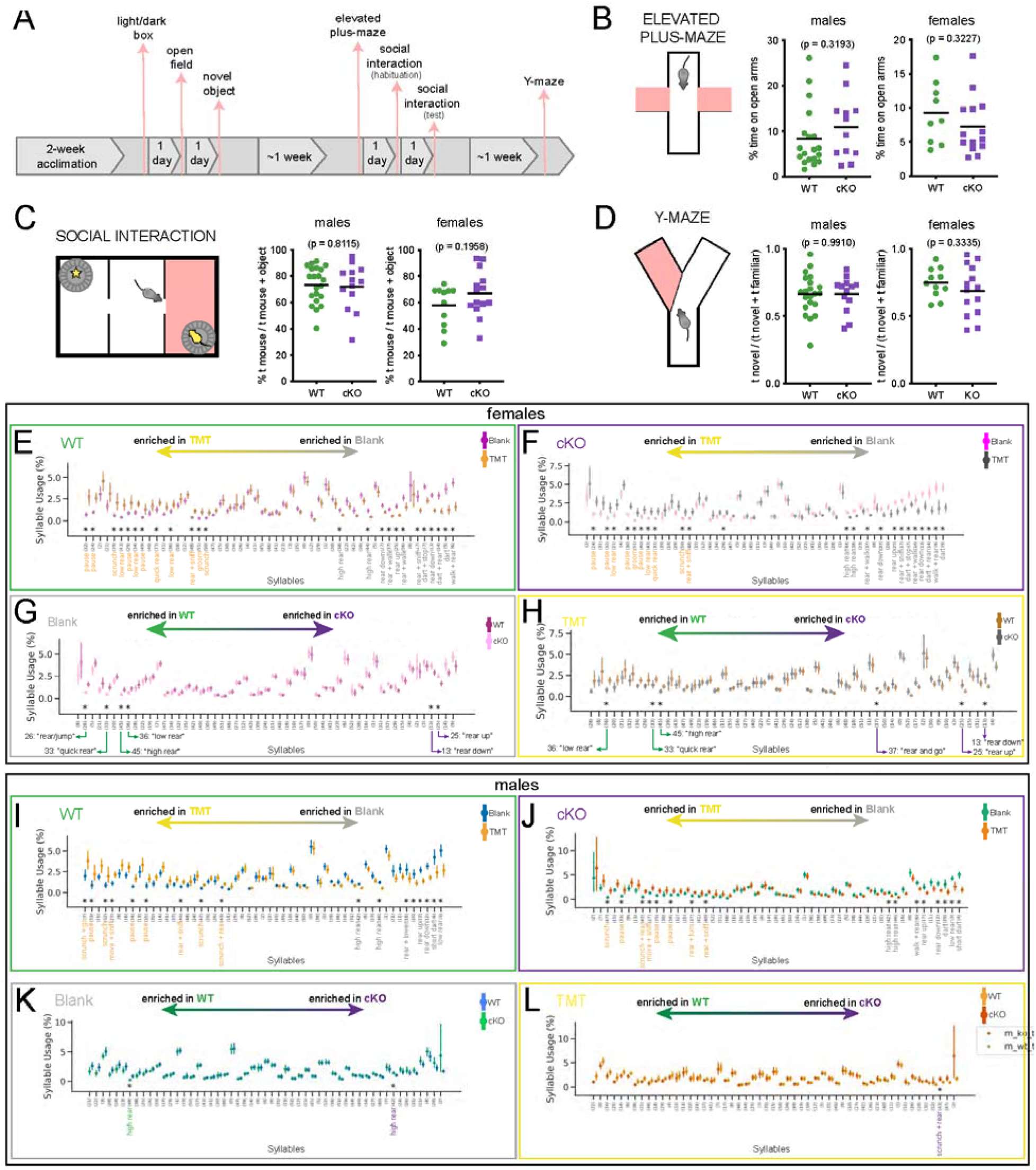
related to Figure 5. **A)** Overview of the timeline used when performing anxiety-related behavior tests. **B-D)** Left, cartoons illustrating the outline of each behavior test; right, summary of the main anxiety-related readout for males and females. The behavioral tests performed were: **B**, elevated plus-maze; **C**, social interaction; **D**, Y-maze. Unpaired t-tests with Welch’s correction were performed; the p-values are shown above the corresponding compared sets of data: bold typeface indicates statistically significant (p<0.05) differences. **E-L)** Syllable usage in female (**E-H**) and male (**I-L**) WT mice during both parts of the experiment, with syllables most enriched in the corresponding portion of the experiment or the genotype as indicated above each graph. Data points in **E-L** represent the average ± 95% confidence interval of the proportion (in %) of test time spent using the corresponding syllable. Significantly different syllable usage (indicated by asterisks) was determined using a Kruskal-Wallis test, post-hoc Dunn’s two-sided test with permutation, and multiple comparisons correction using the Benjamini-Hochberg procedure with a false discovery rate of 0.05.

**Figure S6.**
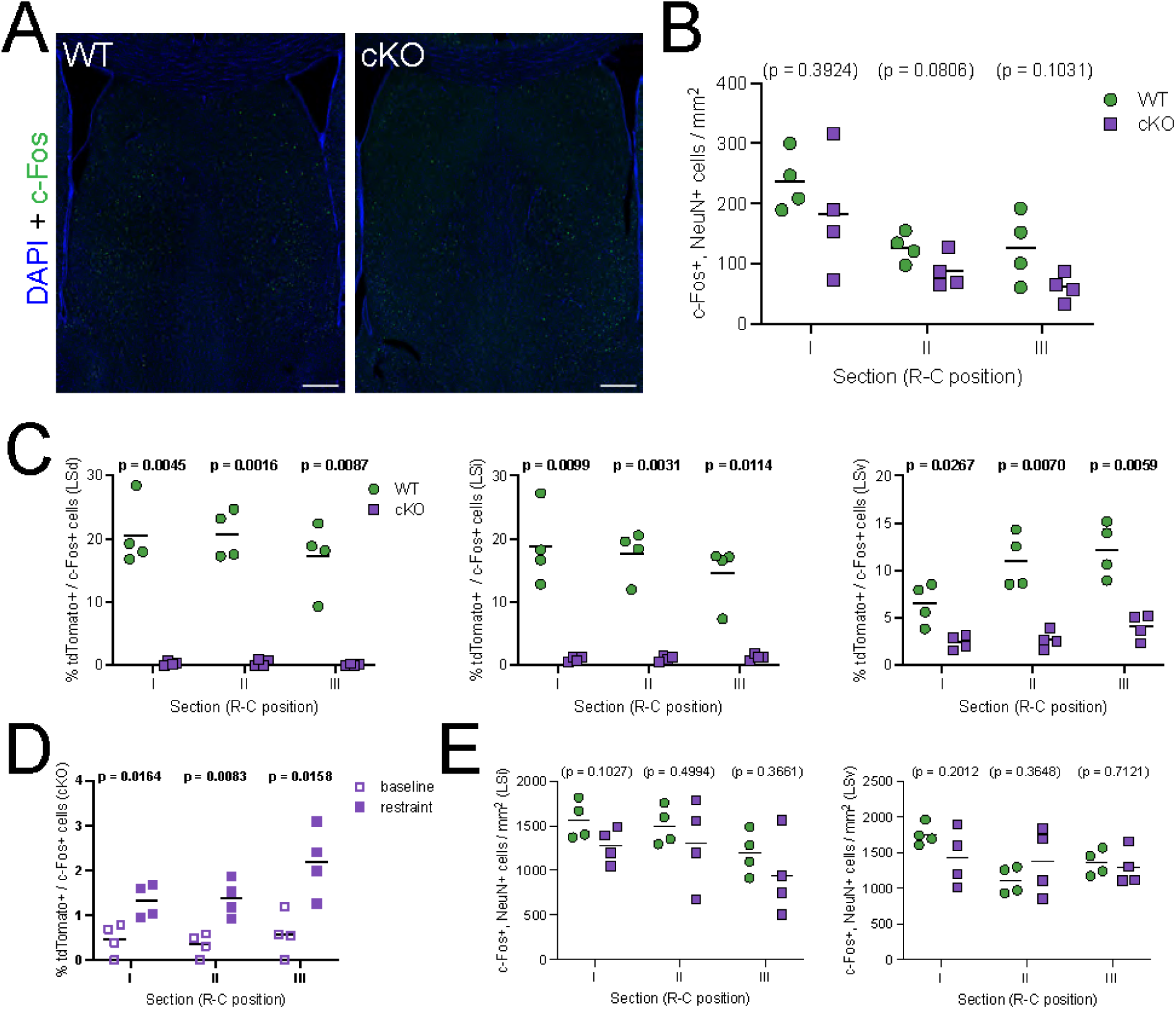
related to Figure 4. **A)** Examples of coronal sections of the septum of WT (left) and cKO (right) age-matched control mice (i.e., not subjected to the forced restraint experiment), immunostained for c-Fos (green) and counterstained with DAPI (blue). **B)** Quantification of the density of c-Fos+ neurons per mm^2^ in the entire LS of WT (green circles, n = 4) and cKO (purple squares, n = 4) control animals. **C)** Proportion of tdTomato+ neurons within the c-Fos+ population in the different subnuclei within the LS of WT (green circles, n = 4) and cKO (purple squares, n = 4) mice subjected to forced restraint. **D)** Proportion of tdTomato+ neurons within the c-Fos+ population in the LS of cKO mice, comparing untreated controls (‘baseline’, empty squares) and animals subjected to forced restraint (‘restraint’, full squares). **E)** Comparison of the density of c-Fos+ neurons in the LSi (top) and the LSv (bottom) of WT (green circles) and cKO (purple squares) animals subjected to forced restraint. Unpaired t-tests with Welch’s correction were performed; the p-values are shown above the corresponding compared sets of data: bold typeface indicates statistically significant (p<0.05) differences.

